# Audio-visual combination of syllables involves time-sensitive dynamics following from fusion failure

**DOI:** 10.1101/771634

**Authors:** Sophie Bouton, Jaime Delgado-Saa, Itsaso Olasagasti, Anne-Lise Giraud

**Author notes:** Correspondence and requests for materials should be addressed to ALG. Sophie Bouton and Jaime Delgado-Saa contributed equally to this work.

## Abstract

In face-to-face communication, audio-visual (AV) stimuli can be fused, combined or perceived as mismatching. While the left superior temporal sulcus (STS) is admittedly the locus of AV integration, the process leading to *combination* is unknown. Analysing behaviour and time-/source-resolved human MEG data, we show that while *fusion* and *combination* both involve early detection of AV physical features discrepancy in the STS, *combination* is associated in with activity of AV asynchrony-sensitive regions (auditory and inferior frontal cortices). Based on dynamic causal modelling, and neural signal decoding, we further show that AV speech integration outcome primarily depends on whether the STS can quickly converge onto an existing multimodal syllable representation, and that *combination* results from subsequent temporal processing, presumably re-ordering, of discrepant AV stimuli.

## Introduction

Screen-based communication poses specific challenges to our brain for integrating audiovisual (AV) disparities due to either asynchronies between audio and visual signals (e.g. facetime, skype) or to mismatching physical features (dubbed movies). To make sense of discrepant audio-visual speech stimuli, our brain mostly focuses on the auditory input, which is taken as ground truth, and tries to discard the disturbing visual one. In some specific cases, however, AV discrepancy goes unnoticed and the auditory and visual inputs are implicitly *fused* into a percept that corresponds to none of them. More interestingly perhaps, discrepant AV stimuli can also be *combined* into a composite percept where simultaneous sensory inputs are perceived sequentially. These two distinct outcomes can experimentally be obtained using “McGurk effect”^1^, where an auditory /aba/ dubbed onto a facial display articulating /aga/ elicits the perception of a fused syllable /ada/, while an auditory /aga/ dubbed onto a visual /aba/ typically leads to a mix of the combined syllables /abga/ or /agba/. What determines whether AV stimuli are going to be fused^2–4^ or combined^5^, and the underlying neural dynamics of such a perceptual divergence is not known yet.

Audio-visual speech integration draws on a number of processing steps distributed over several cortical regions, including auditory and visual cortices, the left posterior temporal cortex, and also higher-level language regions of the left prefrontal^6–9^ and anterior temporal cortex^10, 11^. In this distributed network, the left superior temporal sulcus (STS) plays a central role in integrating visual and auditory inputs from the visual motion area (mediotemporal cortex, MT) and the auditory cortex (AC)^12–17^. The STS is characterized by relatively smooth temporal integration properties making it resilient to the natural asynchrony between auditory and visual speech inputs, i.e. the fact that orofacial speech movements often start before the sounds they produce^4, 18, 19^. Although the STS responds better when auditory and visual speech are perfectly synchronous^20, 21^, its activity can cope with large temporal discrepancies, reflecting a broad temporal window of integration in the order of the syllable length (up to ∼260 ms)^22^. This large integration window can even be pathologically stretched to about 1s in subjects suffering from autism spectrum disorder^23^. Yet, the detection of shorter temporal AV asynchronies is possible and takes place in other brain regions, in particular in the dorsal premotor area and the inferior frontal gyrus^24–27^. The STS and the IFG regions hence exhibit different functions in AV speech integration, owing to their different temporal integration properties^28^. Interestingly, a relative resilience to asynchrony could confer the STS a specific sensitivity to the incongruence of *physical* features across A and V modalities. A key function of the STS could hence be to resolve AV speech feature discrepancies^29^ via a process requiring a double sensitivity to canonical visual motion (lip movements) and auditory spectrotemporal (formant transitions) cues. On the contrary, the frontal cortex has been widely associated with temporal information processing in both short- and long-term memory^30, 31^. When processing AV speech, the frontal lobe could monitor and estimate the AV temporal sequence^26^.

To characterize the mechanism(s) underlying integration of A and V physical speech features (in the STS), we previously developed a generative predictive coding model^32^ that probed whether cross-modal predictions and prediction errors could be utilized to combine speech stimuli into different perceptual solutions, corresponding to *fused*, i.e., /ada/, or *combined*, i.e., /abga/, percepts (supp. Note 1). The model showed that considering the temporal patterns in a 2^nd^ acoustic formant/lip aperture two-dimensional (2D) feature space is sufficient to qualitatively reproduce participants’ behaviour for *fused*^15, 33^ but also *combined* responses^32^. Simulations indicated that fusion is possible, and even expected, when the physical features of the A and V stimulus, represented by the 2^nd^ formant and lip in the model, are located in the neighbourhood of an existing 2D syllable representation. This is the case for the canonical McGurk stimulus, which falls in the /ada/ neighbourhood, when the input corresponds to the /aga/ visual features and /aba/ auditory features. Conversely, audio-visual stimuli having no valid syllable representation in their 2^nd^ formant/lip neighbourhood (Figure 1A) lead to the sensation that the two (quasi) simultaneous consonants /b/ and /g/ are being pronounced sequentially, e.g. the *combination* percepts /abga/ or /agba/^34–36^. Given the resilience to asynchrony of the STS, the *combination* process likely involves additional brain regions that are sensitive to AV timing.

**Figure 1.**
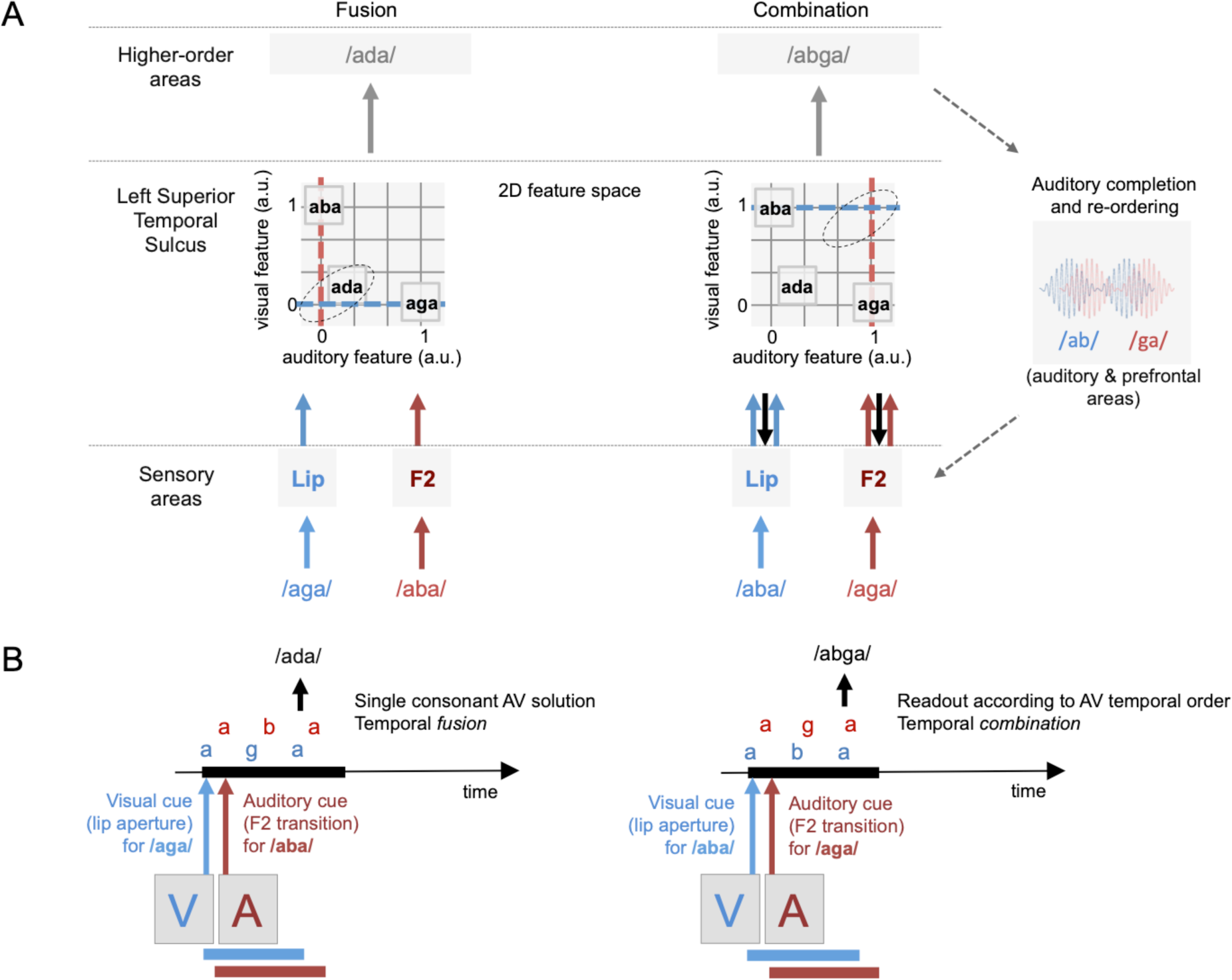
(A) Proposed neurophysiological mechanisms for *fusion* versus *combination*. We posit that after being processed by primary auditory and motion sensitive areas (bottom row), AV inputs converge in the left Superior Temporal Sulcus (STS, middle row) that works as a multidimensional feature space, here reduced to a simple 2D-space in which lip-motion and 2^nd^ speech formant are the main dimensions. The STS is relatively insensitive to AV asynchrony (as depicted in B), but encodes both physical inputs in the 2D-space, converging on the most likely cause of a common speech source given these inputs. In the visual /aga/ - auditory /aba/ condition, coordinates in the 2D space fall close to those of the existing syllable /ada/, which is picked as solution such that the subject senses no conflict. In the visual /aba/ - auditory /aga/ condition, the absence of existing /aXa/ solution at coordinates crossing triggers post-hoc reconstruction of the most likely cause of the inputs via a complex consonant transition /abga/ (true temporal sequence), with occasional temporal inversions of the sound sequence /agba/^35^. Both *combination* outputs require additional interaction with time sensitive (prefrontal and auditory) brain regions. Grey arrows represent the STS output as readout by higher order areas. Blue and red arrows represent visual and auditory inputs, respectively. (B). Discrepant audio (A) and visual (V) syllabic speech units /aXa/ are represented within a critical time-window for integrating them as a single item coming from the same source. The auditory percept is either a McGurk *fusion* /ada/ (left) or a *combination* percept /abga/ (right).

These theoretical data lend support to the recent proposal that the neural processes underlying AV *combination* differ from those involved when AV stimuli are *congruent* and for AV *fusion*^5, 37^. In this study, we address whether AV *combination* involves extra time sensitive processes, and attempt to dissociate between two alternative hypotheses: 1) whether *combination* readily involves fine detection of AV asynchrony (outside the STS, most likely in the left IFG) and the percept arises on-line from the real order with which each phoneme is detected (Figure 1B, right panel), or 2) whether a retrospect reconstruction of the most plausible AV sequence is triggered by the impossibility for AV stimuli to converge on a single 2D (lip/formant) representation in the STS. The latter hypothesis has two experimental implications that we sought to put at test. The first one is that the delay to converge on a plausible solution should be longer in *combination* than *fusion*, because *combination* requires solving AV discrepancy by explicitly serializing auditory and visual inputs into a complex consonant transition (Figure 1B). Extra processing time for *combination* relative to *fusion* should manifest both in reaction times and in the timing of neural events. The second implication is that *combination* should emerge from early AV comparison in the STS and only subsequently involve auditory and (articulatory) prefrontal cortices, to produce an ordered and articulable new composite syllable. In this process, the STS is expected to have a pivotal role and we should therefore observe enhanced functional interactions between the STS and these additional brain regions during *combination* relative to *fusion*.

## Results

To address whether *combining* AV stimuli results from on-line sensitivity to AV asynchrony or from *fusion* failure in the STS, we used the two canonical McGurk conditions, which give rise to either the *fused* percept ‘ada’ or ‘ata’, or a *combined* solution ‘abga’, ‘agba’, or ‘apka’, ‘akpa’. Although these stimuli are artificial (see ^38–40^ for ecologically valid audiovisual stimulation), they allow for a rigorous parameterization of the various AV speech integration outcomes.

We first ran two behavioural experiments carried out in distinct groups of participants. Both experiments involved vowel-consonant-vowel syllables of the type aXa denoted /aXa/ for audio and [aXa] for visual. These AV stimuli were used across three different conditions: (i) a *congruent* condition in which auditory and visual inputs corresponded to the same syllable (stimuli /ada/ + [ada] and stimuli /ata/ + [ata]), and two *incongruent* conditions in which auditory and visual inputs could give rise to either (ii) a *fusion* percept (stimuli /aba/ + [aga] and stimuli /apa/ + [aka]) or (iii) a *combination* percept (stimuli /aga/ + [aba] and stimuli /aka/ + [apa]) (Figure 2C). All stimuli were video clips showing either a female or a male articulating aXa stimuli belonging to the phoneme family ‘bdg’ or the phoneme family ‘ptk’. In a first experiment, 20 participants performed a repetition task. Instructions were the same as those given in the McGurk & MacDonald article (1976): participants watched the videos and were asked to repeat what they “heard” as fast as possible (Figure 2A), with no restrictions on the pronounced syllable. Detailed behavioural analyses are presented in the Methods section.

**Figure 2.**
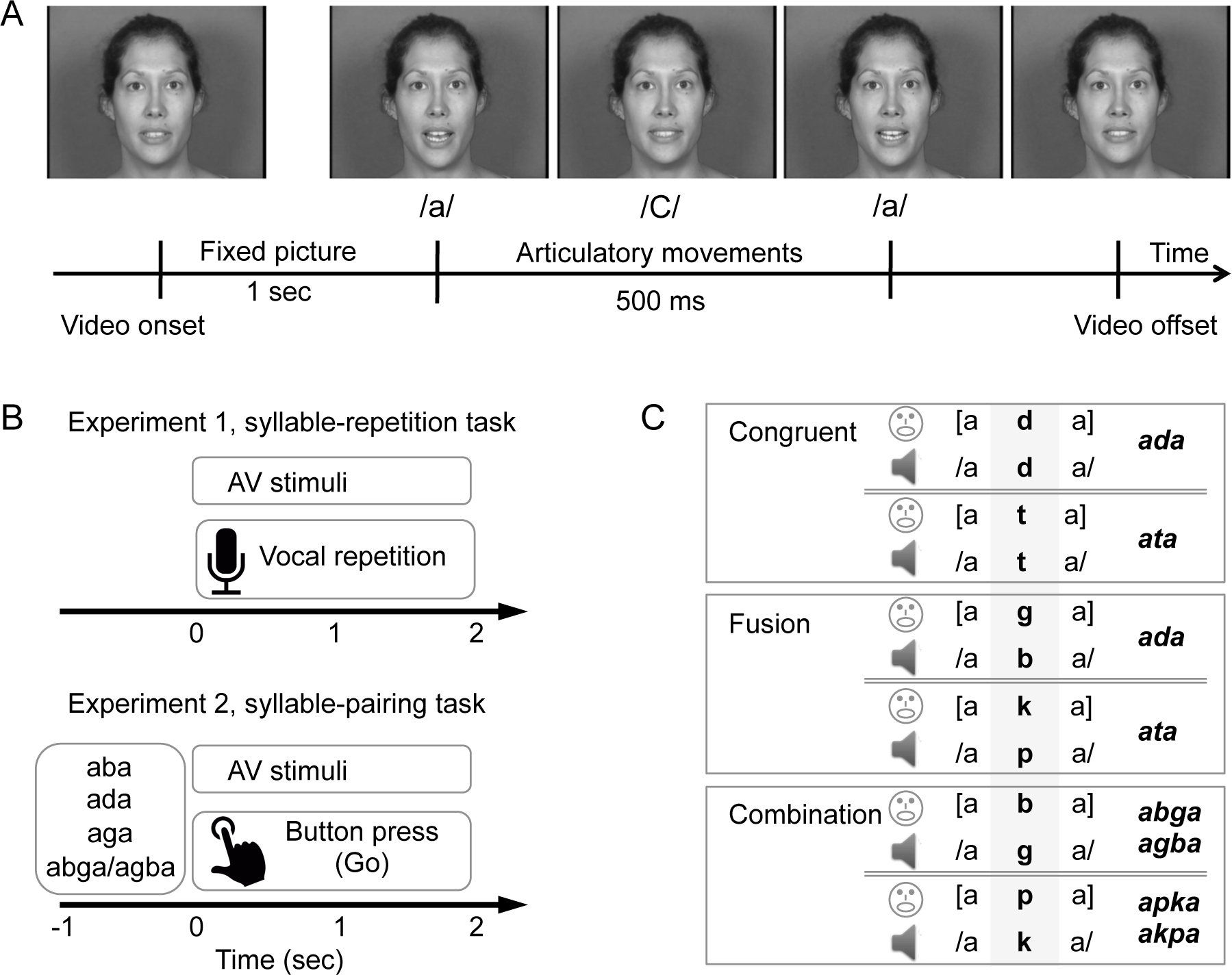
(A) Typical time course of audio-visual stimuli. (B) Example trials from experiments 1 and 2. Experiment 1: trials started with a 1s fixation period, followed by a videoclip showing a speaker pronouncing a syllable. Participants had to repeat the syllable they “heard” as fast as possible. Experiment 2: trials started with a 1s written syllable, followed by a short videoclip showing a speaker pronouncing a syllable. Participants were instructed to press a button as fast as possible if the written syllable matched the syllable they perceived from the AV videoclip. (C) Experimental conditions used in the behavioural and MEG experiments. The same three conditions, labelled ‘*congruent*’, ‘*fusion*’, and ‘*combination*’ were used in the behavioural and neuroimaging experiments. In each condition, stimuli combined a video track and an audio track, and used two consonant families: either ‘bdg’ or ‘ptk’. Right panel: the answer of interest (expected response depending on the exact AV stimulus *combination*) is shown in bold-italic.

### AV *combination* takes longer than *fusion*

We compared the task’s dependent variables (response of interest rate and response time) between the three conditions (*congruent*, *fusion* and *combination*) using two repeated-measures ANOVAs (see Methods, Figure 3, Table S1, Table S2). In the repetition task (behavioural experiment 1), the report rate for each response of interest (i.e. the percentage of ‘ada’ and ‘ata’ responses in the *fusion and congruent* conditions, and the percentage of ‘abga’, ‘apka’, ‘agba’ and ‘akpa’ responses in the *combination* condition) differed across conditions (*F*(2,38) = 275.51, *P* < .001, partial η^2^ = 0.68), irrespective of the consonant family (*F* < 1) or speaker gender (*F* < 1) (Table S2). Subjects reported more ‘ada’ and ‘ata’ responses in the *congruent* than *fusion* condition (*t*(19) = 8.45, *P* < .001, *Cohen d* = 0.76) (Figure 3A, left panel), but the mean rates for each response of interest were not different across *fusion* and *combination* conditions (*t*(19) = 0.69, *P* > .20, *Cohen d* = 0.03). As expected, in the *fusion* condition participants mostly reported fused and auditory-driven responses. In the *combination* condition they reported mostly combined responses, but also auditory and visually-driven responses (Table S2). In the three conditions, only response times (RTs) associated with the responses of interest were analysed, i.e., ‘ada’ and ‘ata’ responses in the *congruent* and *fusion* conditions, and ‘abga’-‘apka’-‘agba’-‘akpa’ responses in the *combination* condition. RTs differed between conditions (*F*(2,38) = 6.92, *P* = .001, partial η^2^ = 0.65): the syllable heard in the *fusion* condition was repeated as fast as in the *congruent* condition (*t* < 1, Cohen *d* = 0.002), whereas the delay to repeat the syllables heard was longer in the *combination* condition than in the *congruent* and *fusion* conditions (*t*(19) = 3,98, *P* < .001, Cohen *d* = 0.61, difference *combination* – *congruent* = 198 *ms*; *t*(19) = 3.78, *P* < .001, Cohen *d* = 0.59 difference *combination* – *fusion* = 191 *ms*, respectively) (Figure 3A, right panel), irrespective of the consonant family (*F* < 1) or speaker gender (*F* < 1) (Table S2). Thus, RTs indicate that subjects were slower to integrate mismatching audio-visual inputs when they elicited a combined rather than a fused, percept. Importantly, repetition time for ‘ada’ or ‘ata’ was equal whether it arose from congruent or incongruent syllables, showing that AV incongruence was not at the origin of the slower RT for *combination*.

**Figure 3.**
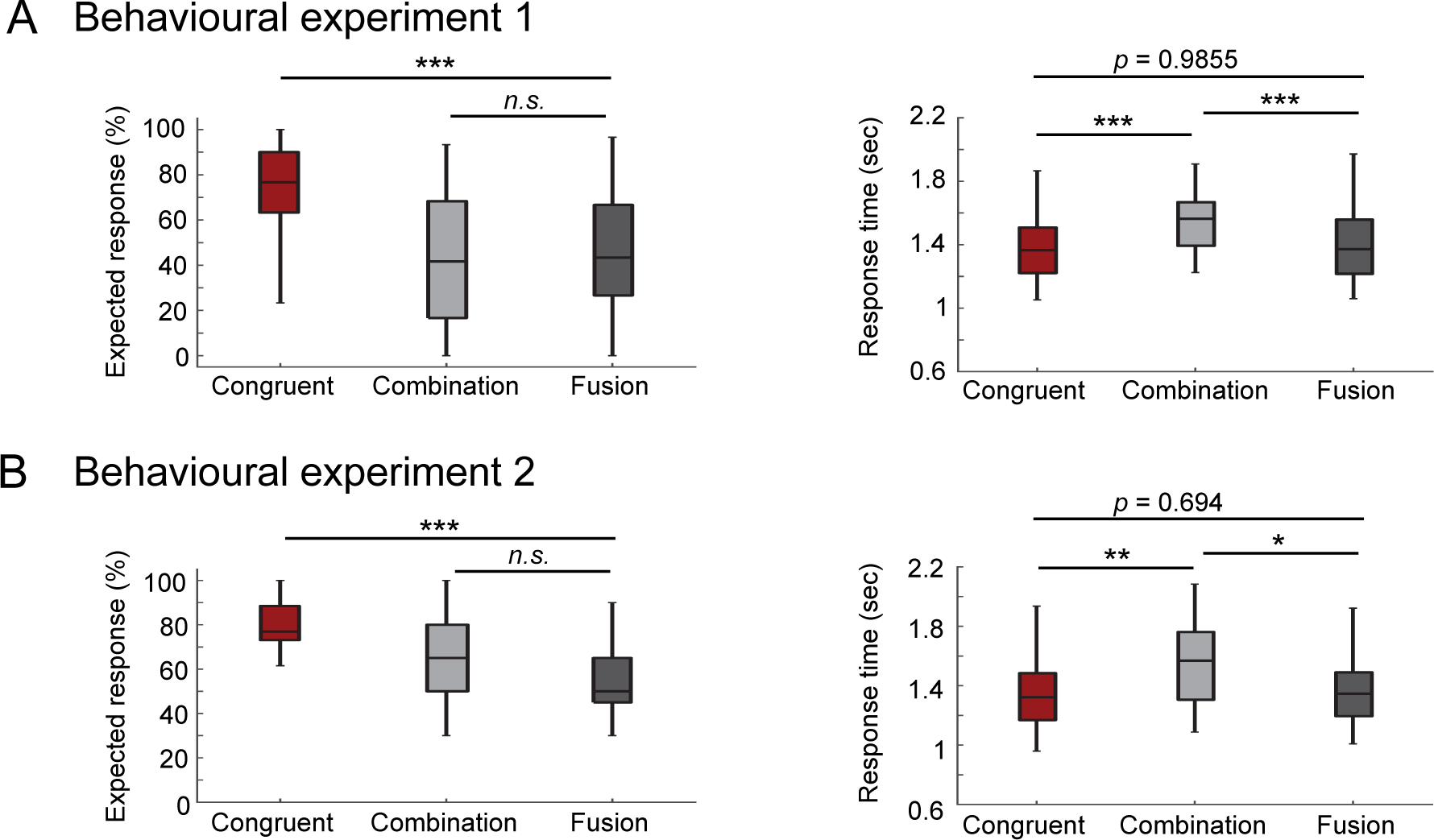
(A) Dependent variables from behavioural experiment 1, syllable-repetition task. (B). Dependent variables from behavioural experiment 2, syllable-pairing task. (A & B, left panels) Rate (%) of responses of interest in each condition, i.e., ‘ada’ or ‘ata’ responses in the *congruent* and *fusion* conditions, ‘abga’, ‘agba’, ‘apka’ or ‘akpa’ responses in the *combination* condition. (A & B, right panels) Response time for responses of interest in each AV condition. In (A & B) error bars correspond to s.t.d. Three stars indicate a significant difference at *P* < .001, two stars indicate a significant difference at *P* < .01, and n.s. indicate a non-significant difference (*P* > .05).

Although, this first experiment supports the second hypothesis that participants should be faster to fuse than to combine AV stimuli, the effect could be biased by the difficulty to plan and articulate a more complex (double consonant) *combination* than a (single consonant) *fusion* syllable. To address this potential issue, we ran a second experiment in which 16 new participants performed a pairing task (behavioural experiment 2), where each trial included a written syllable followed by a video clip. Participants had to identify whether a syllable displayed on the screen before the video matched the syllable that was subsequently heard (Figure 2B). Interestingly, the score for each responses of interest in each condition was similar to that of the repetition task (Figure 3B, left panel). Subjects were better at identifying a *congruent* than an *incongruent* stimulus (*F*(2,30) = 30.26, *P* < .001, partial η^2^ = 0.27). In addition, participants matched the written syllable with the video faster in the *congruent* and *fusion* conditions than in the *combination* condition (*F*(2,30) = 6,84, *P* < .001, partial η^2^ = 0.92; difference *combination* – *congruent* = 206 ms, *t*(15) = 3.06, *P* = 0.08, *Cohen* d = 0.58; difference *combination* – *fusion* = 186 *ms, t*(15) = 2.82, *P* = 0.018, *Cohen* d = 0.54; difference *fusion* – *congruent* = 20 ms, *t*(15) = 0.39, *P* = 0.694, *Cohen* d = 0.08), confirming our previous findings (Figure 3B, right panel), and showing that the extra delay for *combination* does not lie in added articulatory complexity. These data overall suggest that AV discrepancy was more easily solved in the *fusion* than in the *combination* condition, and that the integration of incongruent AV stimuli presumably relies on different neuronal processes depending on whether individuals end-up fusing or combining conflicting AV inputs.

### Global brain dynamics of AV *combination*

To investigate the neural underpinnings of AV *combination*, we recorded brain activity during perception of congruent and incongruent AV stimuli using magnetoencephalography (MEG). Participants watched videos showing a speaker pronouncing a syllable and reported which syllable they heard among 5 alternatives (i.e., ‘aba’, ‘ada’, ‘aga’, ‘abga’, ‘agba’ in the ‘bdg’ family, and ‘apa’, ‘ata’, ‘aka’, ‘apka’, ‘akpa’ in the ‘ptk’ family). Subject’s responses were purposely delayed to avoid temporal overlap between perceptual/decisional processes and motor effects due to button press. Response times hence do not constitute relevant data here, and we only consider the report rates (Figure 4), which were calculated for each condition including congruent responses (i.e., /ada/ or /ata/) in the *congruent* condition, fused responses (i.e., ‘ada’ or ‘ata’) in the *fusion* condition, and combined responses (either VA, i.e., ‘abga’ or ‘apka’, or AV, i.e., ‘agba’ or ‘akpa’) in the *combination* condition. To address whether the components of the AV integration brain network were primarily sensitive to asynchrony or to AV physical features (formants and lip motion), and how these two variables contribute to *combination* versus *fusion*, we also varied the delay between audio and visual syllables. We used 12 different stimulus onset asynchronies over a temporal window ranging from −120 ms audio lead to 320 ms audio lag (40 ms step), a window corresponding to a range in which *fusion* responses are expected to dominate over the auditory driven responses^4^; hence maximizing *fusion* reports (Figure 4). This range of asynchrony was chosen not to disrupt the integration process, as within this temporal window A and V inputs are perceived as simultaneous^22, 41^. We can hence probe AV asynchrony sensitivity independent from any behavioural change. In this task, the response of interest rate differed between conditions (*F*(2,30) = 15.99, *P* < .001), whatever the consonant family (*F*(1,15) = 1.98, *P* = 0.16), speaker gender (*F* < 1) or asynchrony (F<1) (see Table S3 for detailed statistics).

**Figure 4.**
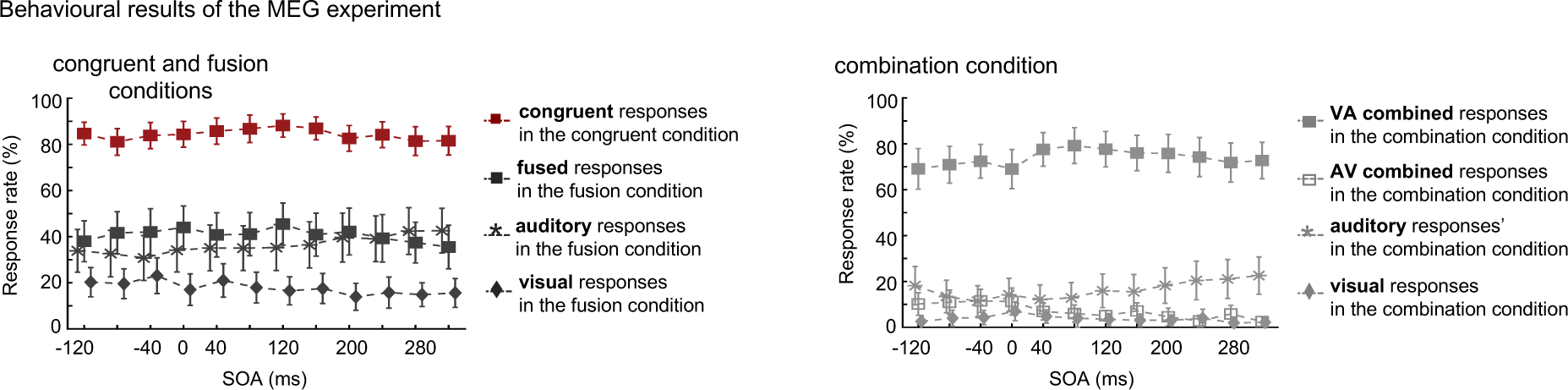
MEG experiment, behavioural results. Response Rate depending on the stimulus onset asynchrony (SOA) between visual and auditory stimuli. A negative stimulus onset asynchrony indicates that the auditory leads the visual input whereas a positive stimulus onset asynchrony shows that auditory lags visual. Error bars correspond to s.e.m. The left panel shows responses in *congruent* and *fusion* conditions: response of interest rates in *fusion* and *congruent* conditions (filled squares), auditory response rate in the *fusion* condition (dark grey stars), and visual response rate in the *fusion* condition (dark grey diamonds). The right panel shows responses in the *combination* condition only: response of interest rate, i.e., VA (visual-auditory) combined responses (light grey filled squares) and AV (audio-visual) combined responses (light grey squares) in the *combination* condition, auditory response rate in the *combination* condition (light grey stars), and visual response rate in the *combination* condition (light grey diamonds).

After checking the global validity of the results in sensor space (Fig S1), we used dynamic source modelling of MEG data to explore the global dynamics of AV *combination* relative to *fusion* and *congruence* conditions (Fig S3). We analysed the evoked activity in six regions of interest showing the strongest incongruence effect (incongruent > congruent): namely the left PAC (Primary Auditory Cortex), MT (Middle temporal visual area), STS (Superior Temporal Sulcus), STG (Superior Temporal Gyrus), IFG (Inferior Frontal Gyrus) and ATC (Anterior Temporal Cortex) (see Fig S2 for a spatial location of the corresponding scouts). Figure 5 shows basic contrasts of conditions for the STS and IFG (see Fig S4 for results in all regions). We found a common statistical effect of *combination vs. congruent* and *fusion vs. congruent* in the STS at ∼100 ms pre-auditory stimulus onset (Figure 5, blue and red), indicating that the STS could signal upcoming AV incongruence, presumably based on visual information (^29, 42, 43^, see also next data analyses). This was the only time point and location where *fusion* and *combination* had a similar response pattern. Importantly, *combination* was associated with a denser activity pattern than *fusion* in both the IFG and the STS. *Combination* gave rise to a pre-audio activity increase in the IFG at −80ms, followed by activity increases in the IFG at +120ms, in the STS at +350ms and again in the IFG at +750 ms. Figure 5 additionally shows a more transient effect in left superior temporal areas (∼350 ms post-auditory stimulus onset). *Combination* was also associated with specific neural activity decreases at ∼200ms in the left STS, and ∼350 ms in the left IFG, which are not readily interpretable using direct comparisons across conditions. These first results provide a global picture of the event sequence at play in our experimental paradigm, and although all events cannot be explained by simple contrasts, the analysis overall shows that *combination* is associated with enhanced processing in both the STS and the IFG.

**Figure 5.**
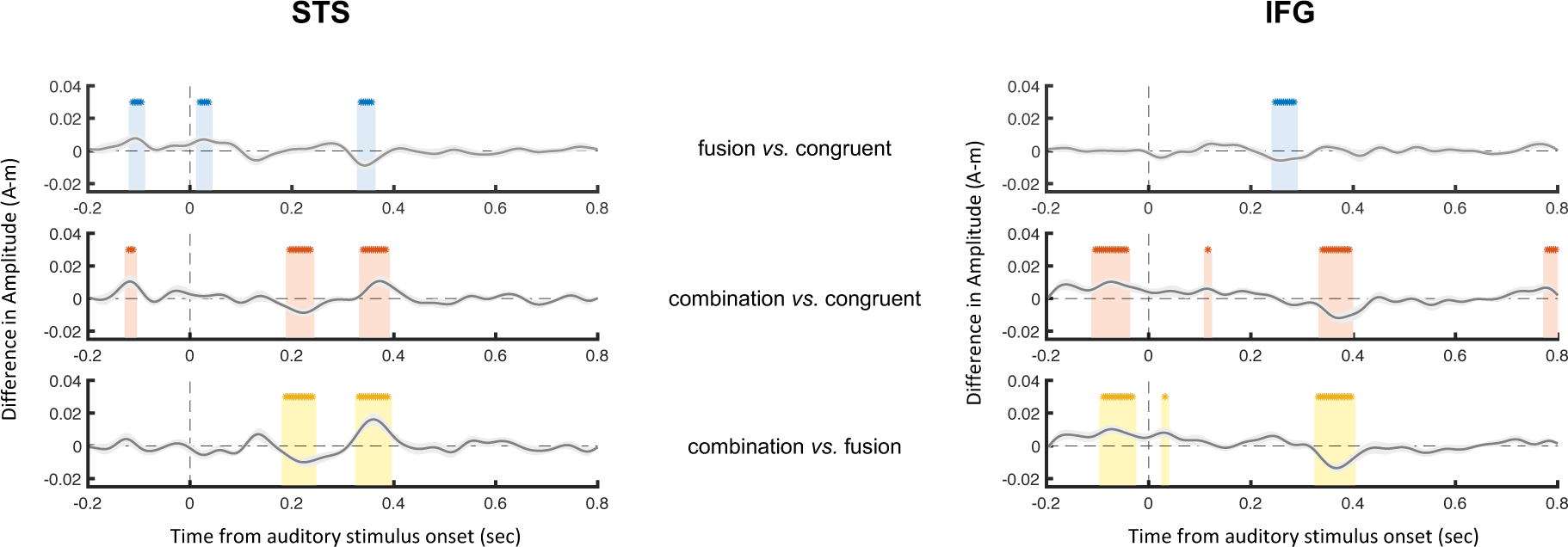
Differences in event-related activity between conditions, in the two regions of interest, i.e. the STS and the IFG (*fusion* > *congruent* conditions in blue, *combination* > *congruent* conditions in red, *combination* > *fusion* conditions in yellow). Stars indicate significant Student *t-test values that* were estimated in each difference: *fusion vs. congruent* conditions in blue, *combination vs. congruent* conditions in red, *combination vs. fusion* conditions in yellow (*P* < 0.05, corrected for multiple comparisons using FDR).

### Directional connectivity patterns for *Fusion* vs. *Combination*

The behavioural results and the dynamic source modelling MEG data both indicate that AV *combination* was more demanding than *fusion*, suggesting that combining discrepant stimuli might involve extra resources and perhaps a different neural network than fusing them. Based on previous modelling work^32^ we conjectured that the left STS plays a pivotal function, i.e. that the involvement of extra neural resources for *combination* could be triggered by the impossibility to converge locally on a bimodal syllable solution. To further explore this hypothesis, we first probed directional functional coupling across the 6 previously defined regions of interest of the left hemisphere (ATC, IFG, STS, STG, MT, and PAC) using dynamic causal modelling (DCM). This analysis was not time-resolved, thus only showed dominant connectivity patterns throughout the experimental trials. We found that *fusion* and *combination* had radically different neural dynamics, characterized by a dominant modulation of feed-forward and feedback connectivity from and to the STS, respectively (Figure 6). *Fusion* was associated with increased connectivity from the STS toward ATC and MT, and a decrease from MT to STS, consistent with the propagation of the *fusion* solution to higher-order regions and the update of visual motion representation as a function of the solution elaborated in the STS. In line with previous studies showing that visual speech prediction could directly influence the activity in auditory cortex^39, 44^, connectivity also increased from MT to PAC and decreased from PAC to ATC during *fusion.* In contrast, *combination* was associated with increased connectivity from IFG and PAC to STS. Finally, connectivity also increased from PAC to both STS and IFG, and decreased from STS to ATC, from ATC to IFG, from STG to PAC.

**Figure 6.**
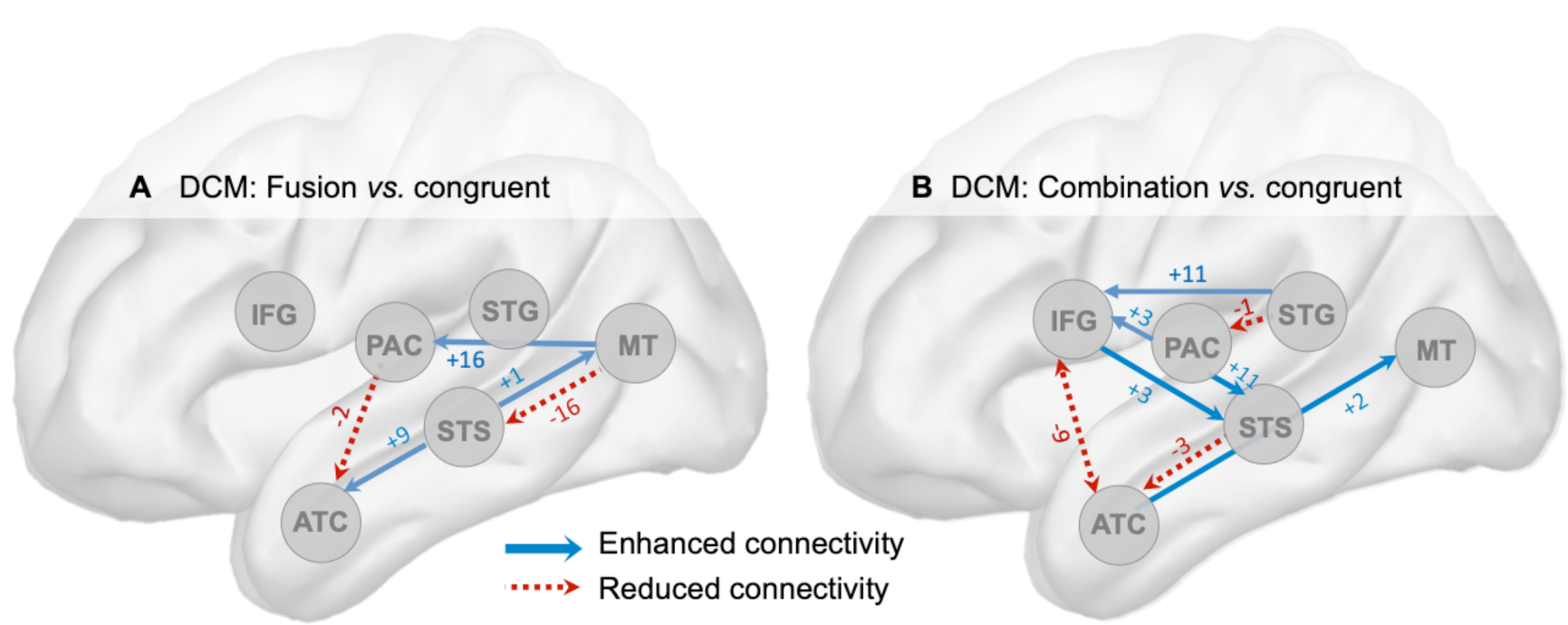
Dynamic causal modelling (DCM) of connectivity using event-related responses across the six main regions involved in audio-visual speech integration. The circles represent the sampled sources: primary auditory cortex (PAC), mediotemporal cortex (MT), the superior temporal gyrus (STG), the left superior temporal sulcus (STS), the inferior frontal gyrus (IFG) and the anterior temporal cortex (ATC). All connections and their values reflect enhanced or reduced connectivity of *fusion* (A) and *combination* (B) responses, relative to responses in the *congruent* condition. We tested the differences between conditions using Parametric Empirical Bayes (PEB) models. Rather than comparing different network architectures, we performed a post-hoc search by pruning away parameters that did not contribute to the model evidence (p<0.05). These results in a sparse graph where the connections shown are those that contributed significantly to the model evidence. Red dotted lines: reduced connectivity; Blue lines: enhanced connectivity.

The functional connectivity results confirm the central role of the STS in both *fusion* and *combination*. They further show that in *fusion*, the STS dispatches information (presumably about the identified syllable) to other brain regions for recognition and sensory representation updating, whereas in *combination*, it centralizes information from higher-order regions, possibly regions that are more sensitive to the precise timing of the AV events than the STS. Crucially, these results show that connectivity from and to the IFG was only enhanced in *combination*.

### Time-sensitivity and neural determinants of *Combination* vs. *Fusion*

As hypothesized the observed behavioural response delay and the specific sensitivity of the IFG for *combination*, may be accounted for by two distinct neural mechanisms. The IFG could directly track the A and V inputs on-line, or it could reconstruct retrospectively the AV order after *fusion* failure in the STS. To address time-sensitivity during AV speech processing and enquire whether the *combination* outcome results from one or more specific neuronal event(s), we used a general linear model (GLM). We regressed auditory input-locked neuronal activity for each of the six regions of interest at each time point against key trial-wise quantities (see Methods for a full description of the GLM): (i) *lip aperture* associated with the stimulus condition with values [1, 0.6, 0.37] for [*Combination*, *Congruent*, *Fusion*] (Fig S5), (ii) *second formant* associated with the stimulus condition with values [0.5, 0.5, 0.2] for [*Combination*, *Congruent*, *Fusion*] (Fig S5), (iii) *AV asynchrony* values (from 0 ms to 320 ms) irrespective of whether auditory or visual signal came first, (iv) *AV physical incongruence* associated with the stimulus, i.e. congruent to incongruent lip motion/2^nd^ formant patterns, (v) *combined output, i.e.* participant’s responses with the value 1 if the AV inputs were combined or 0 if they were not, and (vi) *fused output* with the value 1 when AV inputs were fused and 0 when they were not.

Consistent with the direct comparison of experimental conditions and the DCM analyses, the main effect of AV physical incongruence (Figure 7A) was significant first in the STS during a time period ranging from 150 ms to 70 ms pre-audio onset. In our experimental setting the second input could be predicted from the first one: participants always received /ada/ + [ada] in the congruent condition, or /aba/ + [aga] in the *fusion* condition, or /aga/ + [aba] in the *combination* condition. This high predictability presumably explains that cross-modal effects emerged very early in the STS (Figure 7A and Fig S6), as previously shown^44^. Strong predictions likely resulted from overlearned AV associations and short-term adaptations likely to occur in the current experimental setting^45^. By including the variation of lip aperture across conditions in the GLM, we could confirm that lip-aperture was primarily reflected in the STS activity prior to the audio onset, and that it could hence reflect the prediction of subsequent incongruence (Figure 7A and Fig S6). The interaction between lip aperture and AV incongruence was significant only before auditory onset whereas the interaction between auditory features and AV incongruence was never significant. Together, these results show that AV incongruence may reflect expectation effect before auditory onset but real AV integration process after auditory onset.

**Figure 7.**
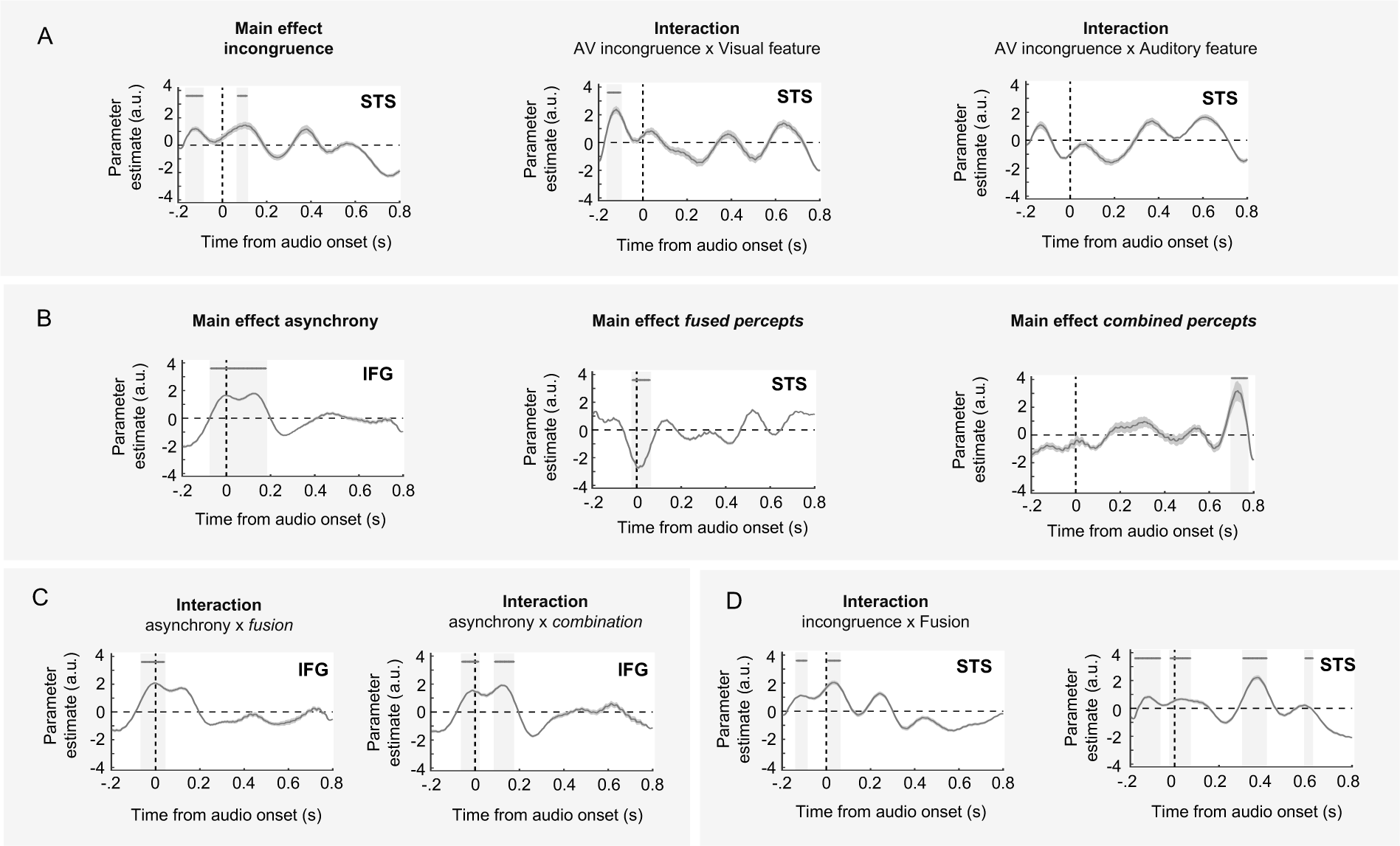
Results of the GLM analysis showing the temporal dynamics of normalized beta in two areas of interest, i.e. the STS and IFG. (A) The correlation between the STS activity and the regressor AV Incongruence emerges significantly before auditory onset at the same time than the STS activity correlates with lip aperture. (B & C) The correlation between IFG and AV Asynchrony peaks around −50 ms, and stops as soon as a fused percept is identified in the STS but last until 200 ms post-auditory stimulus onset when the AV inputs cannot be fused (i.e. in *combination* condition). (D) Combined percept correlates in the IFG at ∼700 ms, after a recursive processing of the AV incongruence in the STS. Thick horizontal lines and light grey areas indicate time windows where parameter estimates diverge significantly from zero at a temporal cluster-wise corrected p-value of 0.05. The shaded error bounds indicate s.t.d. STS. superior temporal sulcus; IFG. inferior frontal gyrus

Confirming the contrast of evoked responses shown in Figure 5, AV physical incongruence also positively correlated with neuronal activity in the STS ∼50 ms after the auditory input (Figure 7A). This effect occurred in parallel with the tracking of variations in the second formant (Fig S6), reflecting a real integration process in the STS rather than mere expectation. Importantly, the incongruence effect stopped at ∼50 ms for *fusion*, but repeated itself at 400 and 600 ms in *combination* (Figure 7D, bottom and Fig S6).

Confirming recent literature, AV asynchrony was reflected primarily in IFG neural activity (20 ms pre- to 200 ms post-auditory stimulus onset), but also in PAC (around 300 ms post-auditory stimulus onset) (Figure 7B and Fig S6). The sequence of positive coefficients in the IFG and then in PAC speaks to the role of the IFG in contextualising the sensory integration process^46, 47^, as AV asynchrony likely directly triggers expectations and possibly associated memory processes^48^. Crucially, in *fusion* the AV asynchrony effect in the IFG quickly dropped (50 ms post-audio), whereas in *combination* it occurred again at 100-200 ms (Figure 7C). The recurrence of incongruence effects in the STS and asynchrony effects in the IFG in *combination* but not *fusion*, are in line with the behavioural findings indicating that *combination* is more time-demanding than *fusion*.

Finally, fused percepts from *fusion* stimuli (Figure 7B) was reflected in the STS activity as soon as the auditory stimulus occurred, presumably signalling a possible match with the visual expectation. This effect was followed by a significant *fusion* effect in the STG 400 ms post-auditory stimulus onset, possibly reflecting that the syllable selected the STS has a phonological validity (Fig S6). In contrast with fused percepts, combined percepts involved activity in the STS, PAC and IFG, all peaking around 700 ms post-auditory stimulus onset (Figure 7D and Fig S6). This late activity indicates that when the AV input did not match a known syllable (no possible *fusion*), the set of regions previously involved in detecting AV physical incongruence (0-100 ms) was reactivated for generating a combined percept (Figure 7D and Fig S6). This late *combination* effect might signal the elaboration of a double consonant combination by first indexing two distinct positions within the multisensory feature space in the STS, and ordering them in a left pre-frontal time-sensitive region for subsequent articulation.

Altogether, these results indicate that the STS is not involved in detecting AV asynchrony, but that it can predict the auditory input on the basis of the visual stimulus, and can quickly detect AV discrepancy whatever the subsequent outcome (*fusion* or *combination*). In case of *fusion*, activity quickly drops in the STS as it converges on a solution, whereas in *combination* AV timing-sensitive activity reoccurs in the IFG along with an incongruence effect in the STS. All PAC, IFG and STS exhibit sustained activity until a complex (consonant sequence) solution is elaborated. Crucially, the sequence of neural events in *combination* is concluded by a late effect in the IFG that does not reflect asynchrony tracking (Figure 7B), but presumably a more abstract timing process, likely retrospect ordering of the AV stimuli.

### Neural decoding of syllable identity for *Fusion* vs. *Combination*

Having established that the STS could quickly detect inconsistencies between auditory and visual physical features, and that the IFG was sensitive to AV asynchrony, we posited that syllable identity of combined syllables (i.e., ‘abga’, ‘agba’, ‘apka’ or ‘akpa’) might be decoded from neural activity of both the IFG and STS, whereas syllable identity of fused syllables (i.e., ‘ada’ or ‘ata’) might only be decoded from the STS.

We hence probed whether neural activity expressed in the STS and IFG held reliable information about syllable identity using two decoding analyses on the trials with responses of interest, to classify (1) ‘abga’, ‘agba’, ‘apka’ and ‘akpa’ responses from *combination* condition vs. ‘ada’ and ‘ata’ responses from congruent condition, (2) ‘abga’, ‘agba’, ‘apka’ and ‘akpa’ responses from *combination* condition vs. ‘ada’ and ‘ata’ responses from *fusion* condition (Figure 8B and Fig S7). We used *time-resolved* decoding to keep track of the information propagation sequence. In line with previous findings^49^, we observed that local evoked activity from one region was sufficiently discriminable to permit syllable categorization using a maximum correlation coefficient classifier (see Methods). We determined whether neural responses related to combined *vs*. congruent percepts could be separated (Figure 8A). The neural data could be classified in the IFG already at ∼360 ms post auditory stimulus onset, and then in the STS at ∼380 ms (Figure 8A). Conversely, neural responses related to combined *vs.* fused percepts could be discriminated only in the STS at ∼170 ms, ∼340 ms and ∼650 ms post auditory stimulus onset (Figure 8B). These analyses confirmed two crucial points: (1) *combination* was the only percept that could be decoded from neuronal activity outside of the STS, confirming the crucial implication of IFG when combining discrepant AV inputs; (2) combined vs. fused syllables elicited discriminable activity of the STS at ∼170 ms, suggesting that from this time point on the impossibility to fuse AV inputs in the STS was susceptible to trigger a sequence of events leading to the identification of a combined syllable.

**Figure 8.**
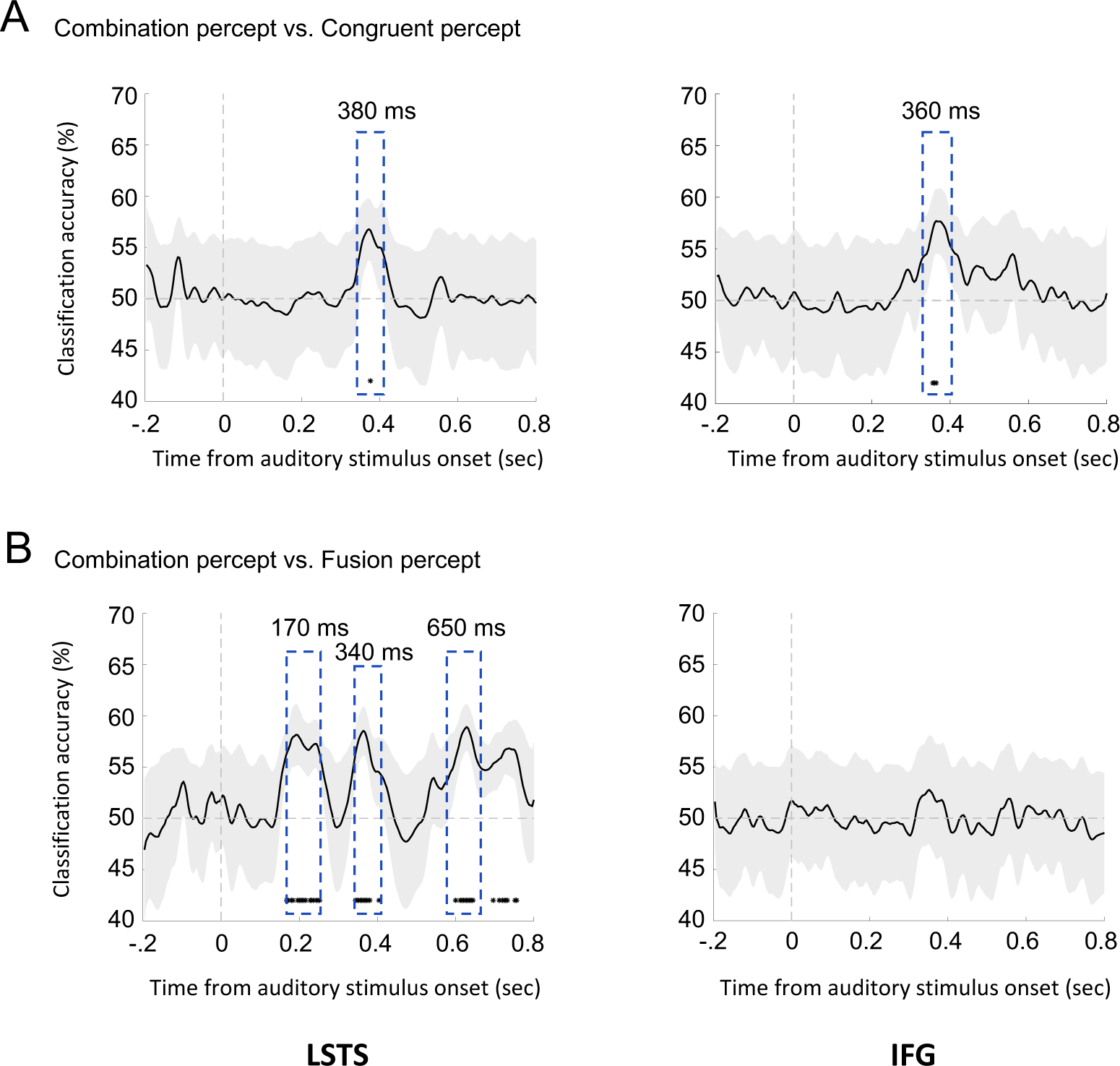
Decoding in the Left Superior Temporal Sulcus (STS) (left panels) and the Inferior Frontal Gyrus (IFG) (right panels). (A) Time course of univariate classification (accuracy) for ‘abga’, ‘agba’, ‘apka’ and ‘akpa’ responses from *combination* vs. ‘ada’ and ‘ata’ responses from *congruent*. Univariate classification between the responses of interest from *congruent* and *combination* was possible in the IFG at ∼360 ms and in the STS at ∼380 ms. (B) Time course of univariate classification (accuracy) for ‘abga’, ‘agba’, ‘apka’ and ‘akpa’ responses from *combination* vs. ‘ada’ and ‘ata’ responses from *fusion*. Univariate classification of responses of interest from *fusion and combination* conditions was possible in the STS at ∼170 ms, ∼340 ms and ∼650 ms, but not in the IFG.

## Discussion

While the mechanisms leading to AV speech *fusion* are relatively well understood, those leading to AV stimulus *combination* are still unknown. Based on a previous computational model, we conjectured that AV *combination* follows from the difficulty to map the auditory and visual physical features in a multisensory space presumably located in the left STS^32^. AV *combination* would hence result in a more demanding processing sequence than AV fusion, possibly involving post-hoc temporal reordering of the auditory and visual input. The model was adapted to McGurk stimuli, and hence worked well with only lip and 2^nd^ formant values (even though more features are presumably at play in AV speech integration). According to this simple 2-dimensional model, *fusion* occurs when the physical features of discordant AV stimuli fall in the vicinity of those corresponding to a lexically plausible and simple (single consonant transition) speech representation, whereas *combination* occurs when the physical features do not find AV matching features (Figure 1A). In this view, after *fusion* failure, *combination* demands additional processing, possibly by frontal areas, to covertly generate a plausible speech sound consistent with the AV input. The alternative scenario would be that *fusion* occurs when AV asynchrony is non-detectable (e.g. because the visual stimulus is a weak predictor), whereas *combination* arises when A and V onsets can clearly be sequentially perceived (e.g. strong visual predictor). The two scenarios lead to distinct predictions regarding the delay with which a *combination* percept arises. In the first case, combining discrepant AV stimuli should take significantly longer than fusing them, i.e. the time needed for *fusion* failure and active generation of alternative operations such as ordering of A and V stimuli. In the second case, the time to fuse and combine AV discrepant stimuli should be about equal, because *combination* mostly depends on the on-line perception of the AV order (Figure 1B).

Our behavioural findings reveal longer delays for reporting AV *combination* than *congruent* and *fusion* percepts, a novel finding suggesting that reporting AV *combination* requires extra processing resources. Although congruent AV speech expected to produce faster and more accurate responses than incongruent AV speech^50, 51^, we did not confirm that AV *fusion* was more time demanding than responding to congruent AV syllables^2, 12, 52–56^. The difference between our results and previous ones is explained by the fact that contrary to previous studies^12, 52–56^, we only analysed trials where subjects effectively experienced *fusion* while discarding failed *fusion* trials (∼45% of the trials).

More importantly, the results of both behavioural studies consistently show that *combination* was more time demanding than *fusion* (+200ms). The second behavioural study even clarified that the effect was not imputable to added articulatory demands. When examining source-resolved MEG responses in the STS, i.e. the key region for integrating audio and visual speech information^17, 57–59^, we found that neural responses for *combination* were already delayed at the earliest stage of AV speech integration, and were hence unlikely to reflect mere attention or effort mechanisms. The response delay for *combination* was hence partly associated with extra processing time in the STS.

To further understand the nature of the processing delay for AV *combination* relative to *fusion*, we explored how much the STS was sensitive to AV asynchrony, and whether other brain regions could be involved in on-line vs. retrospect ordering of A and V stimuli leading to *combination*. Our MEG experimental design hence involved different stimulus onset asynchronies. Although the behavioural effect of AV asynchrony was weak (Figure 4), as expected within the chosen asynchronous range^22, 41^, we found both left PAC and IFG to be sensitive to this parameter, to the point that a classification algorithm from the IFG activity could decode the contrast between *combination* and congruent percepts. This finding presumably reflects the implication of the IFG in perceptual speech tasks requiring precise temporal parsing or sequencing of speech signals^28, 60, 61^.

Unlike the IFG, the STS was insensitive to asynchrony, but highly sensitive to the auditory and visual physical features incongruence^17, 62^, and in line with previous findings, it signalled very early whether the AV stimuli were discrepant, with a first effect driven by visual predictions (i.e., the auditory cortex response induced by visual input), and a second one following the auditory input driven by AV mismatch (Arnal et al., 2009).

### The role of the STS in the *fusion*/*combination* dynamic divergence

Although it is established that the STS integrates the information coming from A and V modalities, it is still not known whether it processes similarly or differently AV stimuli leading to *fusion* or *combination*. Using a GLM approach we found that the STS was the first region to signal physical incongruence between the two sensory modalities. The incongruence effect was even anticipated by the STS based on prediction from the visual stimuli. A possible explanation for this anticipatory effect is that the STS quickly estimates whether to expect a precise auditory input (strong visual prediction) or rather a set of possible auditory inputs (weak visual prediction)^29^. When visual prediction is weak (e.g., visual /aga/), the STS could more easily fuse auditory and visual inputs (within ∼100 ms), whereas when visual prediction is strong (e.g., /aba/), perceived incongruence is potentially stronger, in some cases resulting in a *combination* percept. In other words, AV *fusion* could partly depend on the confidence associated with the expected input^38^.

However, according to our predictive model of AV syllable integration^32^, the most important factor determining *fusion* is whether the two stimuli meet close to an articulatory valid syllable representation within the 2^nd^ acoustic formant/lip aperture 2D space. In the McGurk *fusion* case, visual /aga/ and auditory /aba/ fall in the vicinity of the 2D /ada/ representation, which quickly appears as a valid solution. This scenario is supported by the decoding analyse showing that neural activity in the STS signals the identity of fused syllables 200 ms before that of combined syllables. Importantly, by combining DCM and the “perceptual output” part of the GLM analyses, we show that the incongruence was both registered and solved extremely rapidly in the STS, and that the outcome was propagated forward (to the IFG and ATC). *Fusion* only differed from congruent processing in that the STS output also flowed backward to MT, presumably to update the visual motion model of the *fusion* syllable. The backward STS influence on lower sensory areas (MT)^63, 64^, provides an interesting illustration of how predictive coding could apply to AV integration^32, 65, 66^.

In the *combination* case, since there is no 2D syllable representation at visual /aba/ and auditory /aga/ coordinates, the STS cannot readily converge on a viable articulatory solution. Activity in the STS rises 200 ms post-auditory stimulus onset like *fusion* but remains sustained until 400 ms; during this time lapse, the STS gets recursively involved in a more complex integration process involving PAC, STG and IFG. These findings suggest that *combination* required a tight coordination between the temporal network and the IFG^39^, presumably to organize the temporal serialization of AV inputs.

### The Inferior Frontal Gyrus tracks AV asynchrony

Time perception is an adaptive function involving different brain regions such as the cerebellum, the hippocampus, and the frontal cortex^67, 68^. The specific response of one region rather than another mostly depends on the duration of the stimulus received^69^ and the task performed^70, 71^. While neocerebellar regions have a specific function in the computation of time intervals, the frontal cortex maintains, monitors and organizes temporal representations^24, 72^. Among frontal areas, the dorsolateral prefrontal cortex (DLPFC) can be modulated by the memory load due to stimulus asynchrony, whereas regions traditionally considered to be involved in speech processing, such as the lower frontal cortex and the upper temporal cortex, can be modulated by the temporal complexity of auditory and visual inputs^26^. These findings are consistent with our observation that the early IFG involvement in AV integration does not relate to feature identification, but is specific to AV timing. The IFG tracked AV asynchrony, at least within the range used in the experiment (from −120 ms auditory lead to 320 ms auditory lag), without translating into a misalignment sensation. This confirmed that this range of AV asynchrony is registered while being perceptually well tolerated^4, 22, 41^. Importantly, asynchrony tracking alone did not explain all the combination effects we observed in the IFG, notably it did not account for the latest effect. This suggests that more elaborated/abstract temporal processes also take place in the IFG during *combination*.

Finally, we observed that the asynchrony effect in the IFG was followed by a similar effect in PAC, a finding that fits well with the observation that regularity in tone sequences modulates IFG activity, and that the estimated intervals propagate from the IFG to PAC^46, 47^. Our study confirms that the IFG originates descending signals about estimated precision or sensory stimulus predictability^73^. Temporal order sensitivity in the left IFG might for instance allow listeners to predict word order from syntactic information^74, 75^, a functional role that is compatible with the retrospect sequentialization of AV stimuli.

### On-line AV temporal tracking, versus post-hoc temporal ordering in *combination*

When auditory and visual signals are well aligned, integrating AV syllables is an easy and relatively low-level process, but when they are asynchronous or physically discrepant, integration involves a more complex dynamics. Our findings show that *combination* results both from the identification of auditory and visual features in the STS, and from the detection of fine AV temporal asynchronies in the left IFG. The global information flow directionality (DCM) and the event sequence (GLM) indicate that *fusion* and *combination* arises from a visual pre-activation of the STS, which is either matched by a compatible audio (*fusion*) and further processed as known syllable (*fusion*), or discarded by an incompatible one and further processed as temporally asynchronous AV sequence. Our results hence support that *combination* is triggered by a failure of AV *fusion* in the STS, and reflects *a posteriori* reconstruction of the most plausible AV sequence.

We found no behavioural or neurophysiological arguments for the alternative scenario in which temporal ordering in *combination* occurs on-line. Although the IFG was sensitive a AV asynchrony in a relatively early timeframe, the late *combination* effects are not accounted by on-line temporal tracking. The combination of visual [aba] and auditory /aga/ can take two forms^35^: one (‘abga’, 72% of the responses) that respects the real AV temporal order, and another one (‘agba’, 8% of the responses) that does not (for similar results on the two *combination* percepts formed with /aba/ and [ada], see^36^). The production of an “erroneous” combined percept ‘agba’ strongly supports a post-hoc process resulting from recursive activity across the STS, PAC, STG and IFG, rather than from on-line sequencing, which offers less possibilities of mixing the AV stimuli.

## Conclusion

The present findings contribute to delineate the hitherto unknown mechanisms of AV *combination.* By showing that *combination* percepts arise from a two-step process consisting in first registering AV physical incompatibility in the STS, and then reordering the stimuli by AV asynchrony sensitive regions, these results illustrate the primordial role of recursive/predictive mechanisms in AV speech integration, and unravel the dynamics leading to the elaboration of novel constructs that best explain the cause of AV stimuli.

## Methods

### Subjects

Twenty healthy subjects participated in the first behavioural experiment (9 males - age range: 20-28 years), 16 subjects in the second behavioural experiment (9 males – age range: 21-31 years), and 15 took part in the MEG study (10 males – age range: 21-24 years). The number of subjects in experiment 1 was defined in accordance with previous studies^3, 15, 16, 37, 55, 62, 76–78^. All participants were right-handed, French-native speakers, and had no history of auditory or language disorders. Each behavioural experiment consisted of 1 hour-long session performed in a quiet room while the MEG experiment consisted of 2 sessions lasting 2 hours each. All participants were paid for their participation. Ethics permission was granted by the University Hospital of Geneva in Switzerland for the behavioural experiments (CEREH 13-117), and by the Inserm ethics committee in France (biomedical protocol C07-28) for the MEG experiment. All participants provided written informed consent prior to the experiment.

### Stimuli

We recorded natural speech consisting of a man’s and a woman’s face articulating syllables. These two native French-speaker pronounced the syllables /apa/, /ata/, and /aka/ or the syllables /aba/, /ada/, and /aga/. The two syllable continua vary according to the place of articulation; furthermore syllables are voiceless in one continuum, i.e., /apa/, /ata/ and /aka/, and voiced in the other continuum, i.e., /aba/, /ada/, and /aga/. To preserve the natural variability of speech, we used 10 exemplars of each syllable pronounced. Movies were recorded in a soundproof room into a 720 x 480-pixel movie with a digitization rate of 29.97 frames per s (1 frame = 33.33 ms). Stereo soundtracks were digitized at 44.1 kHz with 16-bits resolution.

We created 3 movie categories, which corresponded to the 3 stimulation conditions. *Congruent* videos corresponded to the initial recorded movie of the syllables /ada/ or /ata/. All videos had the same length and lasted 1000 ms. Using the soundtrack, we homogenized the duration of the stimuli: the vocal burst of the first /a/ and the consonantal burst were aligned across videos. The length of the second vocalic part was slightly variable across stimuli. Incongruent *fusion* pairs were created by dubbing an audio /apa/ or /aba/ onto a video [aka] or [aga], respectively. Audio and video were merged from the same speaker. The new soundtrack (/apa/ or /aba/), was systematically aligned to the initial soundtrack (/aka/ or /aga/) based on the vocalic burst of the first /a/ and on the consonantal burst. Incongruent *combination* pairs were created by dubbing an audio /aka/ or /aga/ onto respectively a video [apa] or [aba], using the same alignment procedure. Auditory and Visual parameters of each condition are shown in Figure 1 and in Figure 2.

### Tasks design

Auditory-visual stimuli were presented using Psychophysics-3 Toolbox and additional custom scripts written for Matlab (The Mathworks, Natick, Massachussetts, version 8.2.0.701). Sounds were presented binaurally at a sampling rate of 44100 Hz and at an auditory level individually set before the task via earphones using an adaptive staircase procedure. For each participant, we determined prior the experiments their auditory perceptual threshold corresponding to 80% categorization accuracy. The estimated sound level was used to transmit the stimuli (mean 30 dB sensation level) during the behavioral experiments (Experiment 1 and experiment 2) and MEG experiment.

### Experiment 1, Repetition task

Participants were individually tested and were instructed to watch each movie and repeat what they heard as fast as possible. We used the same instruction provided by McGurk and MacDonald (1976). Participants were asked to repeat as fast as they can what they heard. We did not limit the possible answers to a limited set of syllables. Nevertheless, note that for the three conditions (*congruent*, *fusion* and *combination*), different responses were expected according to our hypotheses (see Fig S1).

### Experiment 2, Pairing task

Participants were individually tested and were instructed to read the syllable written on the screen, then to watch the movie and to press the space bar as fast as possible when the written syllable matched what they heard. In the b-d-g sessions, participants could read ‘aba’, ‘ada’, ‘aga’, ‘abga’ or ‘agba’. In the p-t-k sessions, participants could read ‘apa’, ‘ata’, ‘aka’, ‘apka’ or ‘akpa’.

In the two behavioural experiments, participants were presented with four blocks, each one containing one speaker gender (female or male voice) and one continuum (b-d-g or p-t-k). Each block presented 90 AV stimuli corresponding to 30 *congruent* stimuli (A[d]V[d] or A[t]V[t]), 30 *fusion* stimuli (A[b]V[g] or A[p]V[k]), and 30 *combination* stimuli (A[g]V[b] or A[k]V[p]), for a total of 360 stimuli per subject. Trials were randomly presented in each block, and blocks were randomly presented across participants.

Participants were sat 1 m from the monitor, and videos were displayed centered on a 17-inch Apple MacBookPro laptop on a black background. Sounds were presented through earphones (sennheiser CX 275).

### MEG experiment

Each continuum (/bdg/ and /ptk/) was delivered to participants in two independent sessions of 360 trials each. Participants were asked to perform an identification task. Each trial comprised one video (randomly chosen among the 3 conditions), followed by a 1s silent gap; then, a response screen with ‘aba’, ‘ada’, ‘aga’, ‘abga’ and ‘agba’ in the b-d-g sessions, and ‘apa’, ‘ata’, ‘aka’, ‘apka’ and ‘akpa’ in the p-t-k sessions, were displayed. Syllables were randomly displayed from right to left on the screen to prevent motor preparation and perseverative responses. During MEG recording, the appearance of the response screen was randomly jittered 100, 300 or 500ms after the silent gap. Participants indicated their response by moving a cursor under the syllables and pressing a key to select the chosen syllable as quickly as possible. Subject’s responses were purposely delayed to avoid temporal overlap between perceptual processes and motor effects due to button press. Response times hence do not constitute relevant data. To limit eye movements, subjects were asked to blink only after giving their motor response. After the response, a jittered delay varying from 3 to 5 s led to the next trial.’

Audiovisual alignment in asynchrony conditions. Audio-visual asynchronies were created by displacing the audio file in 33.33 ms increments (frame unit) with respect to the movie file. This process resulted in the creation of 12 different stimulus onset asynchronies over a temporal window ranging from −120 ms audio lead to 320 ms audio lag (40 ms step).

### MEG recording and preprocessing

Brain signals were recorded using Neuromag Elekta with a total of 306 channels composed of 204 axial gradiometers and 102 magnetometers. Recordings were first preprocessed using signal-space separation through Neuromag software MaxFilter. This allows removing signal coming from outside of the electrode sphere which allows removal of EOG (electrooculography) and ECG (electrocardiography) interference among other sources of noise. Originally, signals were sampled at a rate of 1000Hz and were re-sampled at 250Hz for further preprocessing stages. Before MEG recording, headshape was acquired for each participant using Polhemus. After the MEG session, an individual anatomical MRI was recorded (Tim-Trio, Siemens; 9 min anatomical T1-weighted MP-RAGE, 176 slices, field of view = 256, voxel size = 1 x 1 x 1 mm^3^). MEG data were preprocessed, analyzed and visualized using dataHandler software (wiki.cenir.org/doku.php), the Brainstorm toolbox ^79^ and custom Matlab scripts.

### Analysis

#### Experiment 1, Repetition task

We recorded the participant’s vocal response using a microphone. No feedback was provided after each response. The response time was measured as the interval between video onset and start of the syllable repetition from the audio recording on each trial. We also assessed the identification choice made by participants, i.e., the syllable repeated, on each trial.

#### Experiment 2, Pairing task

The number of matching between the written syllable and the video that led to a response of interest served as the measure of syllable identification. The response time was measured as the interval between video onset and the button press on each trial.

In the two behavioural experiments, percentage of responses of interest and response onset latency were calculated for each condition. Percentage of responses of interest ([ada-ata] for *congruent* and *fusion* conditions, or [abga-apka-agba-akpa] for the *combination* condition) was averaged separately across Consonant Family (‘bdg’ and ‘ptk’), Speaker Gender (male and female), and Conditions (*congruent*, *combination* and *fusion*) factors. Response onset latency was calculated and averaged based on responses of interest across each condition). We reported the percentage of congruent responses in the *congruent* condition (i.e. /ada/ or /ata/ responses), the percentage of visual (i.e., /aga/ or /aka/), auditory (i.e., /aba/ or /apa/) and fused (i.e., /ada/ or /ata/) responses in the *fusion* condition, the percentage of visual (i.e., /aba/ or /apa/), auditory (i.e., /aga/ or /aka/), VA combined (i.e., /abga/ or /apka/) and AV combined (i.e., /agba/ or /akpa/) responses in the *combination* condition.

*Behavioural analyses:* Analysis of variance. Percentage of responses was analysed within each experiment (experiments 1 and 2) using a 3 × 2 repeated-measures ANOVAs with Conditions (*congruent*, *fusion*, *combination*) and responses of interest ([ada-ata] responses for the *congruent* and *fusion* conditions, and [abga-apka-agba-akpa] responses for the *combination* condition) as within-subjects factors. For the mean response latency, we measured the interval between video and vocal response onsets, for each type of response of interest. A 3 × 1 repeated measures ANOVA was performed on response times (RTs) with Conditions (*congruent*, *combination* and *fusion*) as a within-subjects factor. All ANOVAs modelled the variables Speaker Gender (female and male), and Consonant Family (‘bdg’ and ‘ptk’) as fixed-factors so as to generalize the results obtained to each speaker and each consonant family tested.

*MEG processing*: Using structural data, brain models for each subject were build using Brain Visa Software ^80^. Individual brain models were mapped to the ICBM-112 brain model template for group-level analysis. Data analysis was performed with Brainstorm ^79^, which is documented and freely available for download online under the GNU general public license. The data considered (trials) started 0.2s before the auditory event and until 0.8s after the auditory input (i.e., the first vowel /a/ in /aXa/). We only analysed trials without eye artefacts or jumps in the signal. We did not conduct the analyses at sensor level as the contribution from different sources is mixed and our aim was to describe the brain network involved in AV speech perception. We computed forward models using overlapping-sphere method, and source imaging using weighted minimum norm estimates (wMNEs) onto preprocessed data, all with using default Brainstorm parameters. A classical solution to the forward model is to use an L2 regularized minimum-norm inverse^81^. The L2 regularization parameter was fixed to 0.1 such that the covariance matrix of the data is stabilized by adding to it an identity matrix that is scaled to 10% of the largest eigenvalue. The wMNEs included an empirical estimate of the variance of the noise at each MEG sensor, which brings both magnetometers and gradiometers into the same basic range of units, allowing the source estimation to be proceed with a combined array of 306 sensors (204 planar gradiometers and 102 magnetometers). We also decimated the MEG data to 250 Hz using an IIR low-pass filter (Chebyshev Type I of order 8).

### Regions of Interest (ROI)

We defined six regions of interest in the left hemisphere, using functional localizers at a group-level (Fig S2). We selected all the regions that had the largest M100 (∼110 ms) auditory evoked responses in all conditions across subjects. ROI analyses were carried out by performing ANOVA across subjects in order to ensure that the selected regions are the ones where more activity was found independently of the conditions.

### Evoked Responses

Signals from each region of interest were extracted and analysed. Evoked responses were computed by averaging MEG signals after source reconstruction across trials for each time sample around stimuli, for each subject and each condition (i.e., *fusion*, *combination* and *congruent*) (Fig S3). We then contrasted the conditions by subtracting their respective ERP response, which allowed testing 3 contrasts: *fusion* minus *congruent*, *combination* minus *congruent*, and *combination* minus *fusion*. Differences in the ERP response were detected across conditions by performing *t* tests against 0. FDR corrections for multiple comparisons were applied over the dimensions of interest (i.e., time samples, regions-of-interest and conditions), using the Benjamini–Hochberg procedure.

### Dynamic Causal Modelling (DCM) analysis procedure

Analysis of functional connectivity through Dynamic Causal Modeling was performed to determine significant changes in connectivity strength across conditions. We performed this analysis using the data from the 6 regions of interest defined above. We modelled forward and backward connections between nodes without pre-specified information, i.e. connections between every pair of nodes were probed. Cross-trial effects were estimated using the congruent condition as a reference, hence modelling all changes in connections that are necessary to explain *fusion* and *combination* conditions. The number of modes for data selection was set to 8 and 1 DCT (Discrete cosine transform) component was used per mode. Given that the signals are in the source space, we set the spatial model parameter to LFP in SPM. The statistical tests for connectivity strength changes were implemented using Parametrical Empirical Bayes method through SPM, creating nested models with different connections switched off each time, and comparing the model evidence obtained. This allows determining which connections affect significantly the predictive power of the model.

### General Linear Model procedure

We then created one GLM and applied this GLM to each time point and each ROI separately. This analysis was performed on single-trial event-related activity, on the trials with responses of interest. The GLM included the following parametric modulators: (i) *syllable family* (ptk or bdg) with values [-1, 1], (ii) *gender of the speaker* in the video (female, male) with values [-1, 1]; (iii-v) *fusion*, *combination* and *congruent* conditions using a matrix with values [0,1] for each column; (vi) *lip aperture* associated with the stimulus condition with values [1, 0.6, 0.37] for [*Combination*, *Congruent*, *Fusion*] (Fig S5), (vii) *second formant* associated with the stimulus condition with values [0.5, 0.5, 0.2] for [*Combination*, *Congruent*, *Fusion*] (Fig S5), (viii) the *AV asynchrony* values that vary from 0 ms to 320 ms, without taking into account whether the auditory or the visual signal came first, (ix) the *AV physical incongruence* associated with the stimulus that is either congruent or incongruent, (x) combined output, i.e. participant’s responses with the value 1 if the AV inputs have been combined (i.e., [abga-apka-agba-akpa]) or 0 if they have not been combined, and (xi) fused output (i.e., [ada-ata]) with the value 1, or 0 if the AV inputs have not been fused. We regressed single-trial MEG signals against these 3 parametric quantities at successive time points from −200 ms to 800 ms following auditory stimulus onset. Obtained time courses for parametric modulators in the GLM were smoothed using bandpass filtering (1-40 Hz) and then averaged across subjects. We next determined the time window where parametric modulators for AV asynchrony, *AV* p*hysical incongruence* and output choice value were significantly different from zero. FDR corrections for multiple comparisons were applied over the dimensions of interest (i.e., time samples, regions-of-interest and number of regressors), using the Benjamini–Hochberg step-up procedure.

### Classification of syllable identity

Decoding analyses were performed with the Neural Decoding Toolbox^82^, using a maximum correlation coefficient classifier on evoked responses in each region of interest. Analysis was constrained to trials with the responses of interest. Two different pattern classifiers were built: one classifier was used to detect the neural activity capable of distinguishing between the response of interest in the *fusion* (i.e., ‘ada’ and ‘ata’ responses) and *combination* conditions (i.e., ‘abga’, ‘agba’, ‘akpa’ and ‘apka’ responses). As the two conditions use incongruent AV inputs, the decoding results reflect only the identity of the outputs. Another classifier was used to detect where neural activity allowed the distinction between the response of interest in the *combination* (i.e., ‘abga’, ‘agba’, ‘akpa’ and ‘apka’ responses) and *congruent* conditions (i.e., ‘ada’ and ‘ata’ responses). Comparing the results of the classifier between fused and combined output with the other classifier between *combination* and *congruent* output allowed us to assess profile similarity between the classifiers. By comparing the curves of the classifiers, we could define the elements that specifically corresponded to the *combination* output.

In the decoding procedure, each classifier was trained to associate MEG data patterns with corresponding stimulus conditions (for each trial, the identity of the syllable perceived). The amount of relevant information in the MEG signal was evaluated by testing the accuracy of the classifier on a separate set of test data. We performed the analyses at each time point, within 1-ms non-overlapping bins.

Decoding analyses were performed on each ROI with a cross-validation procedure where the classifier is trained on a subset of the data, and then the classifier’s performance is evaluated on the held-out test data. For each decoding run, data from the selected trials were divided into sets of 10 trials, and the data from each set of 10 trials were averaged together (see^83^ for a similar procedure). Each decoding run was performed at the group level, pooling all subjects together (see^84^ for a similar procedure). For example, in the first decoding procedure, a pattern classifier was trained to associate MEG patterns with the participants’ responses (the identified syllable, i.e., ‘ada’ vs. ‘abga’) in *fusion* and *combination* conditions. For each decoding analysis, the pattern classifier was trained on the participant’s response. It computed the correlation between MEG data and the syllable identified at each time point, and was trained on 80% of the data, while its performance was assessed on the withheld 20% of the test data. The splitting procedure between training and test data was performed 100 times to reduce the variance in the performance estimate. The classifier computed the correlation between test vectors (i.e., randomly selected mean values of 10 trials in the ROI at each time point) and a vector created from the mean of the training vectors. Each test point took the label of the class of the training data with which it maximally correlated. The reported final classification accuracy is reported as the percentage of correct trials classified in the test set averaged over all cross-validation splits. We then assessed the time window where decision values between the two categories were significantly different from zero (*t* test against zero). FDR corrections for multiple comparisons were applied over the dimensions of interest (i.e., time samples, regions-of-interest and number of classifiers), using the Benjamini–Hochberg step-up procedure.

## Acknowledgments

We thank Jean-Luc Schwartz for support and inspiration. We are grateful to Jean-Luc Schwartz, Alexis Hervais-Adelman, and Valérian Chambon for comments and useful discussions about earlier versions of this manuscript, and Christophe Savariaux for technical support of the video editing. This work was funded by the Swiss National Science Foundation (SNF 320030_149319 and SNF 320030_163040 to A-L.G. and SNF P300P1_167591 to S.B.), and by the Fondation pour l’Audition (RD-2016-5 to S.B.).

## Author contributions

Conceived and designed the experiments: S.B., I.O., A-L.G.; performed the experiments: S.B.; analysed the data: S.B., J.D., I.O., A-L.G.; wrote the paper: S.B., J.D., I.O., A-L.G.

## Data Availability

The behavioural data are available here: https://figshare.com/s/63c773e39a239b72c473 and https://figshare.com/s/0ca3cc102593616cb3b2; The MEG data are available here: https://figshare.com/s/a4ba48347905b0c0790f.

## Supplemental Note 1.

The model builds on three processing levels: a bottom one where two types of sensory units encode key features from each sensory modality (the 2^nd^ acoustic speech formant, and the lip aperture values), an intermediate level where serial units determine the timing and ordering of auditory and visual features, and a top level where recognition units (all possible syllable representations such as ‘aba’, ‘ada’, and ‘aga’) are distributed in a two-dimensional feature space spanned by the speech second formant and lip aperture values. These recognition units, when activated by speech stimuli, generate acoustic and visual predictions at the two lower levels. The three levels are hierarchically organized and all rely on a message-passing scheme that minimizes prediction errors at each level of the processing hierarchy.

---

**Fig S1.**
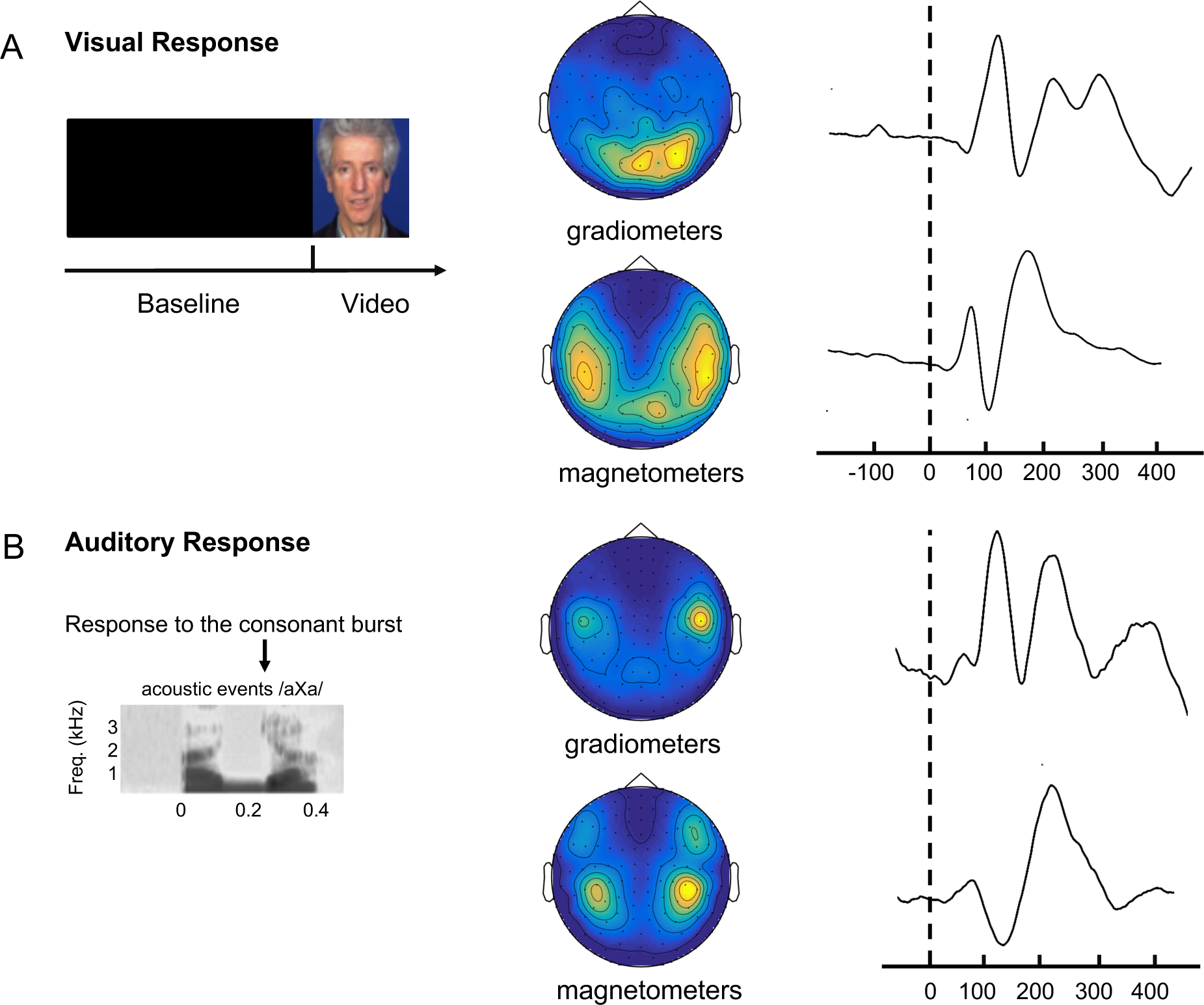
Response profiles on the MEG sensor space. (A) response evoked by visual stimuli, (a video following a black screen) from activity measured with gradiometers (top) and magnetometers (bottom). (B) response evoked by auditory stimuli, i.e. at the consonant burst of the stimuli, from activity measured with gradiometers (top) and magnetometers (bottom).

**Fig S2.**
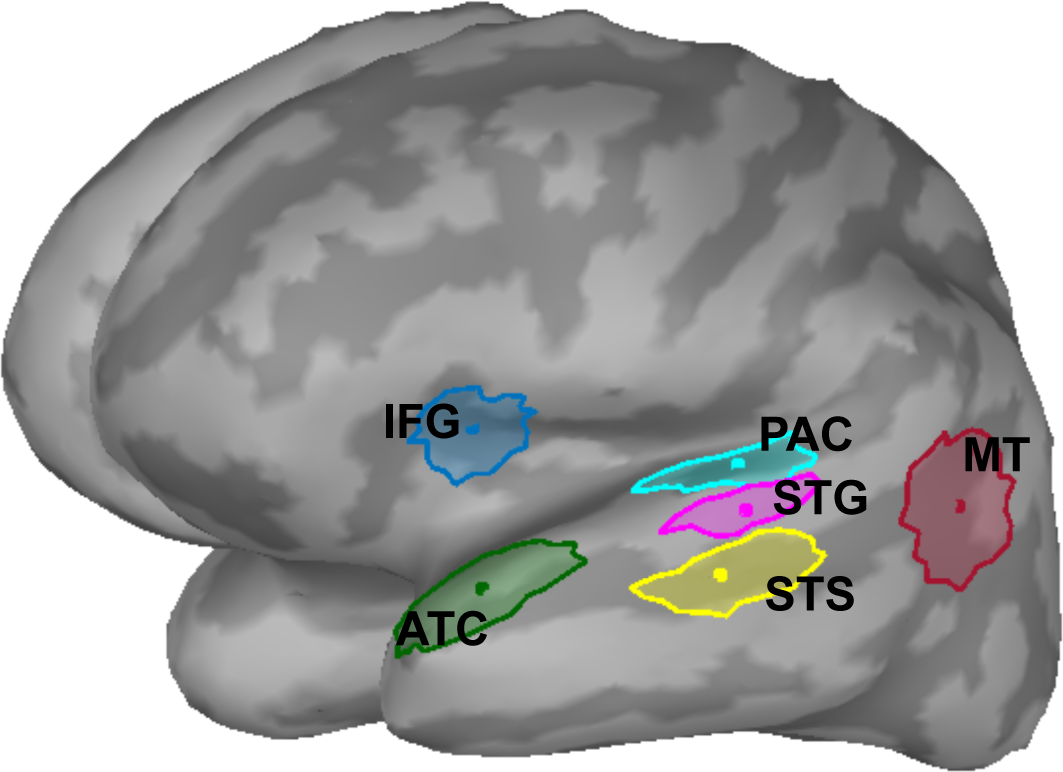
Regions of Interest are the left primary auditory cortex (PAC), the left mediotemporal cortex (MT), the left superior temporal gyrus (STG), the left superior temporal sulcus (STS), the left inferior frontal gyrus (IFG) and the left anterior temporal cortex (ATC).

**Fig S3.**
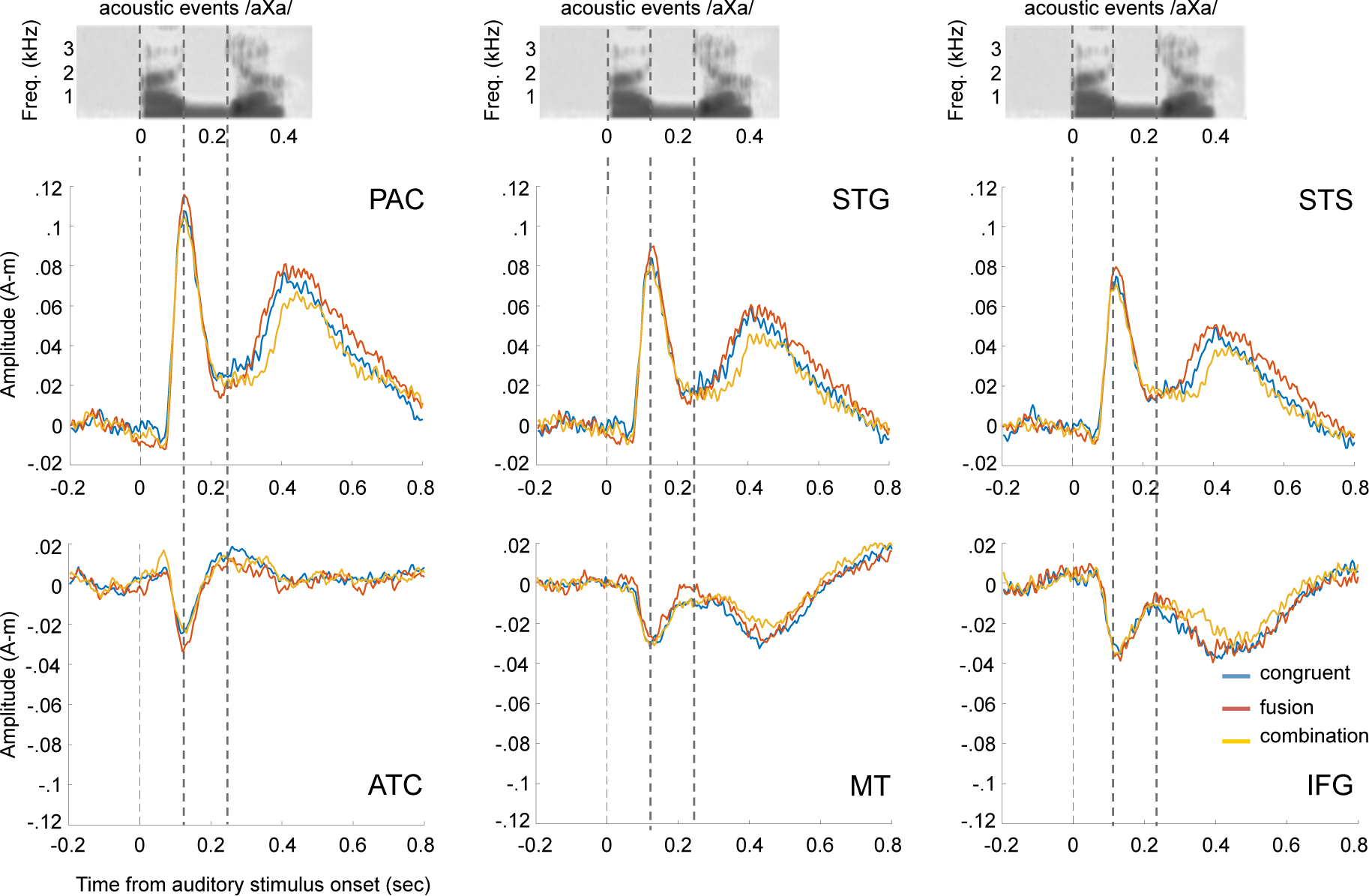
Evoked related field in each condition (congruent in blue, fusion in red, and combination in yellow) in each area of interest. The spectrograms are examples, used to illustrate how neuronal activity is locked to the auditory signal.

**Fig S4.**
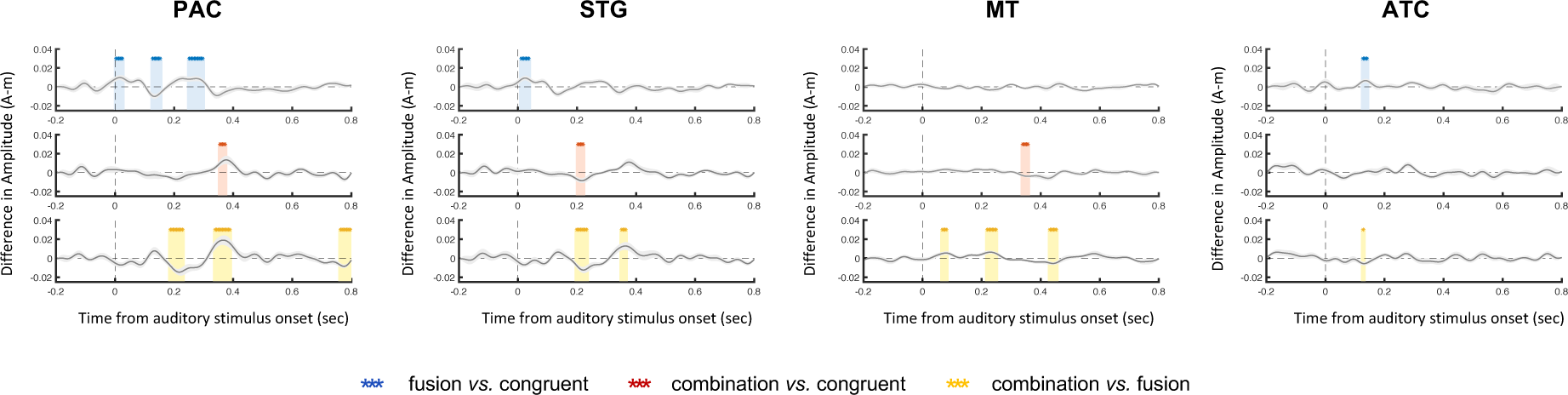
Differences in event-related activity between conditions, in six regions of interest, i.e. PAC, STG, STS, ATC, MT and IFG (fusion > congruent conditions in blue, combination > congruent conditions in red, combination > fusion conditions in yellow). Stars indicate significant Student t-test values that were estimated in each difference: fusion vs. congruent conditions in blue, combination vs. congruent conditions in red, combination vs. fusion conditions in yellow (*P* < 0.05, corrected for multiple comparisons using FDR). PAC. primary auditory cortex; STG. superior temporal gyrus; STS. superior temporal sulcus; ATC. Anterior Temporal Cortex; MT. MedioTemporal Cortex; IFG. inferior frontal gyrus

**Fig S5.**
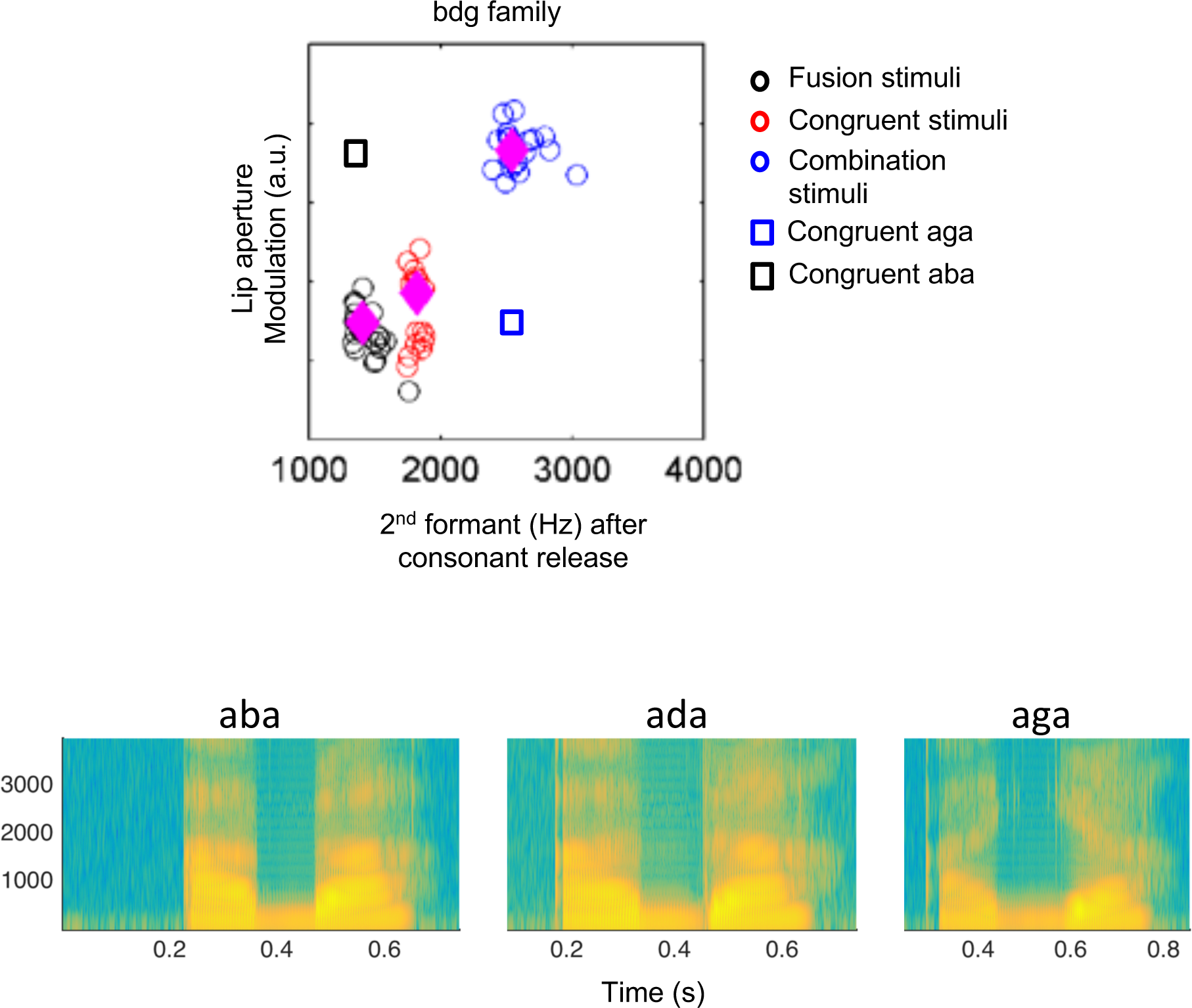
Stimulus features. Top panel: Measure of lip motion amplitude and 2^nd^ formant after consonantal release for each of the 20 stimuli in each stimulus category of the ‘bdg’ family. Diamonds show the median values for each stimulus type. The lip motion amplitude corresponds to the lip aperture difference between maximal aperture at the vowels and the minimal aperture upon consonant occlusion in the middle: ((max_lip aperture 1-min lip aperture)+(max lip aperture 2-min lip aperture))/2. The location of congruent aga and aba stimuli are given for reference, they were not presented to the participants in the study. Bottom panel: spectrogram for sample productions of /aba/, /ada/ and /aga/ sounds.

**Fig S6.**
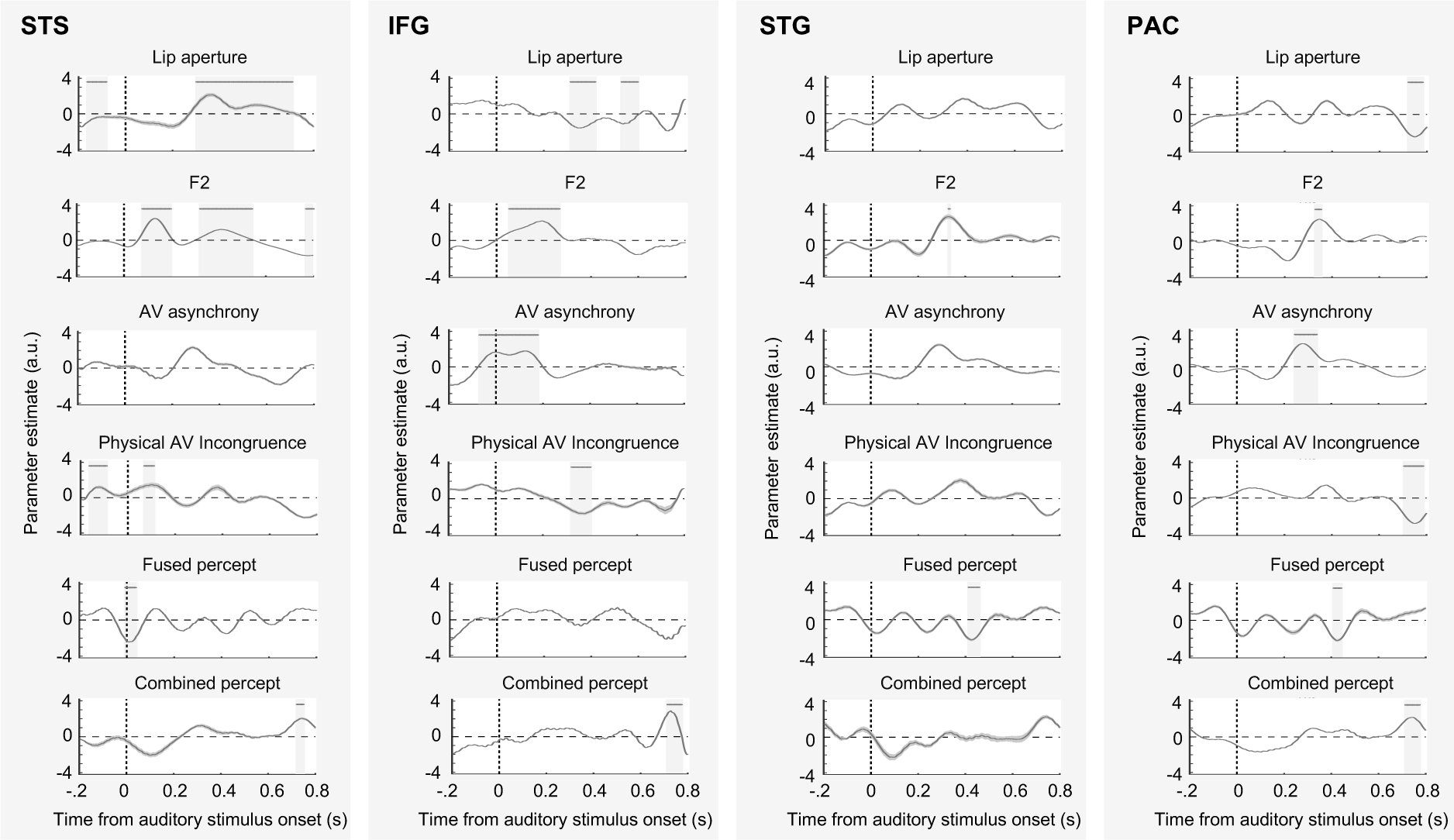
Results of the GLM analyses in four area of interest (STS, IFG, STG and PAC). Thick horizontal lines and light grey areas indicate time windows where parameter estimates diverge significantly from zero at a temporal cluster-wise corrected p-value of 0.05. The shaded error bounds indicate s.t.d. STS. superior temporal sulcus; IFG. inferior frontal gyrus; STG. superior temporal gyrus; PAC. primary auditory cortex.

**Fig S7.**
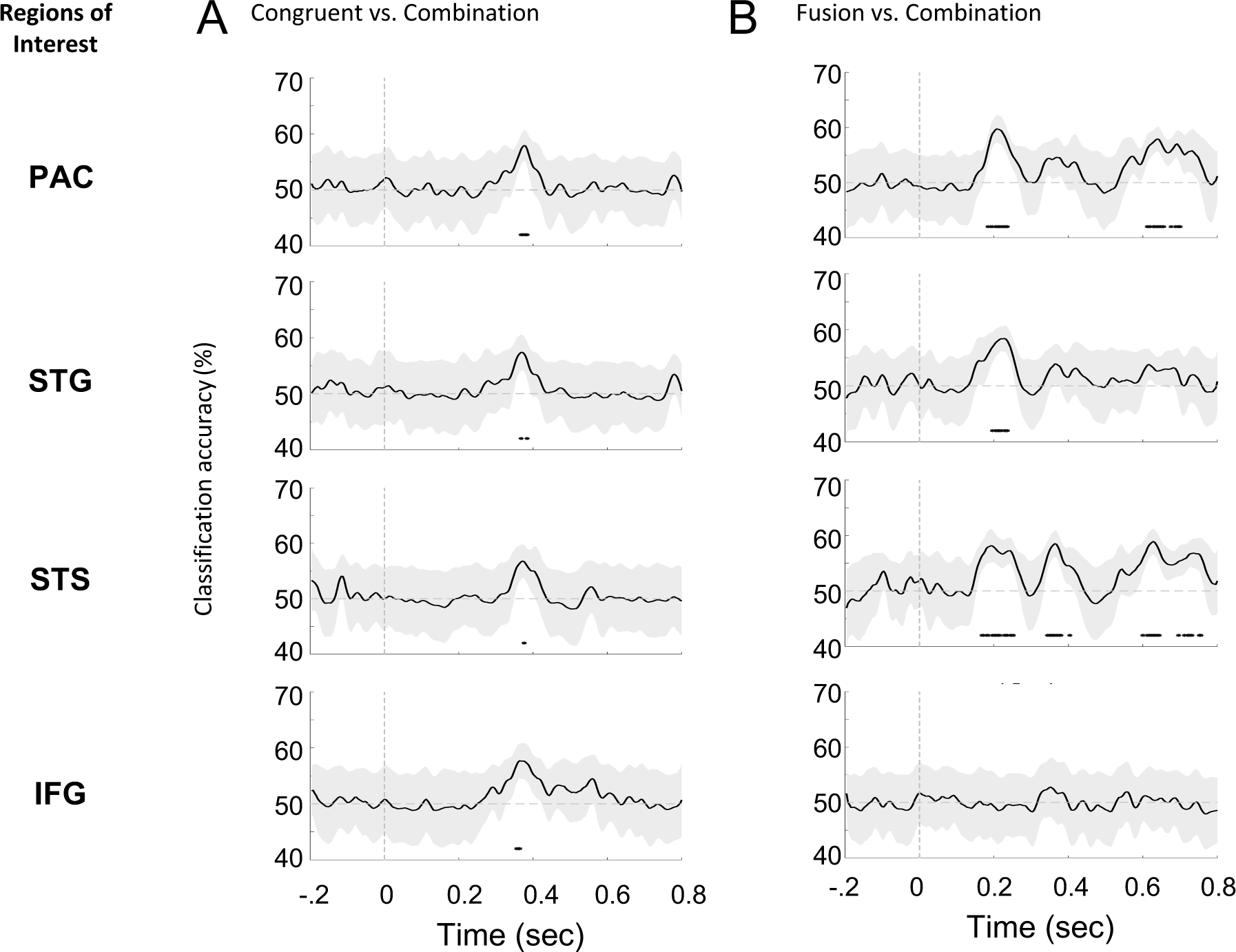
Decoding in the left primary auditory cortex (PAC), the left superior temporal gyrus (STG), the left superior temporal sulcus (STS), the left inferior frontal gyrus (IFG). (A) The time course of the (normalized) univariate classifier results for combined versus congruent percepts. (B). The time course of the (normalized) univariate classifier results for fused percept versus combined percept.

**-- Table S1 --.** Response times and percentage of responses (in brackets) averaged across participants, for each category of syllables produced in experiment 1, according to each condition (congruent, fusion, and combination).

|  |  | Conditions |  |  |
| --- | --- | --- | --- | --- |
|  |  | Congruent | Fusion | Combination |
| Responses | aba / apa | 1.74 (3.2%) | 1.50 (30.9%) | 1.59 (6.0%) |
|  | ada / ata | 1.44 (78.8%) | 1.48 (45.5%) | 1.69 (10.2%) |
|  | aga / aka | 1.62 (10.8%) | 1.55 (12.7%) | 1.74 (21.3%) |
|  | agba / akpa | 1.79 (1.6%) | 1.76 (1.6%) | 1.62 (19.5%) |
|  | abga / apka | 1.77 (1.0%) | 1.66 (1.3%) | 1.58 (24.0%) |

**-- Table S2 --.**
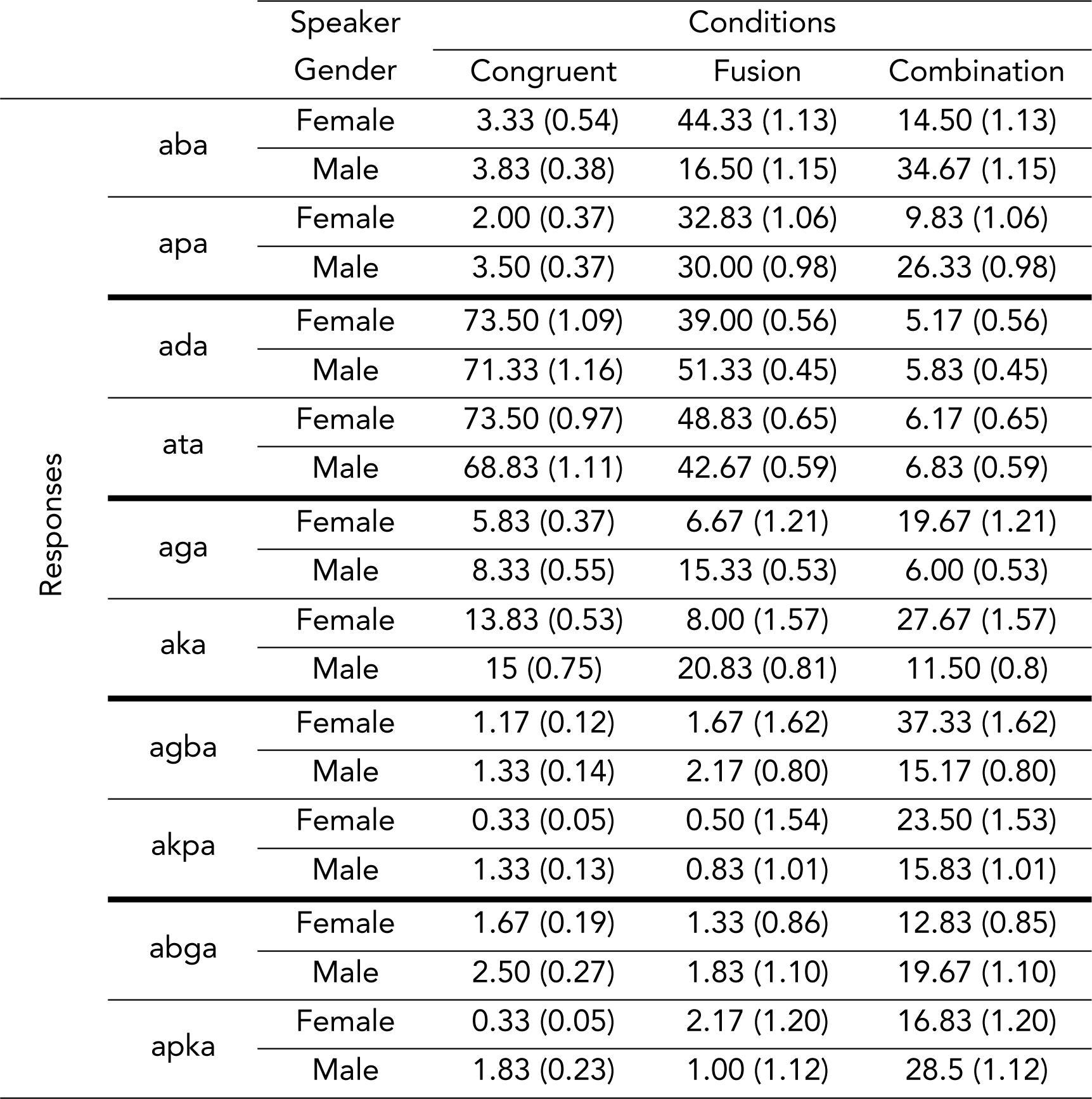
Percentage of responses (mean, and standard deviation in brackets) averaged between participants, for each category of syllables, according to the speaker gender used in the video (male or female) and according to each condition (congruent, fusion, combination), in experiment 1.

**-- Table S3 –.** Behavioural results for the MEG experiment. In the upper part of the table are the results of the ANOVA. In the lower part of the table are the results of the linear regression analysis between each condition and the asynchronies.

| MEG Experiment | d.f. | F | *p* – threshold<br>with *Bonferroni*<br>correction: 0.005 |
| --- | --- | --- | --- |
| Continuum | 1, 14 | 0.98 | 0.34 |
| Voice | 1, 14 | 0.33 | 0.57 |
| Asynchronies | 11, 154 | 0.87 | 0.57 |
| Conditions | 2, 28 | 15.99 | 0.0001* |
| Continuum*Voice | 2, 28 | 0.04 | 0.96 |
| Continuum*Asynchronies | 11, 154 | 0.63 | 0.80 |
| Continuum*Conditions | 2, 28 | 0.99 | 0.38 |
| Voice*Asynchronies | 11, 154 | 1.15 | 0.33 |
| Voice*Conditions | 2, 28 | 0.68 | 0.51 |
| Asynchronies*Conditions | 22, 308 | 1.97 | 0.0063 |
| | $\beta$ | t | p |
| Congruent | -0.003 | -0.519 | 0.604 |
| Combination | 0.020 | 1.815 | 0.070 |
| Fusion | -0.003 | -0.772 | 0.440 |

## References

1. McGurk, H. & MacDonald, J. Hearing lips and seeing voices. Nature 264, 746–748 (1976).

2. Alsius, A., Paré, M. & Munhall, K. G. Forty Years after Hearing Lips and Seeing Voices: The McGurk Effect Revisited. Multisens. Res. 31, 111–144 (2017).

3. Matchin, W., Groulx, K. & Hickok, G. Audiovisual Speech Integration Does Not Rely on the Motor System: Evidence from Articulatory Suppression, the McGurk Effect, and fMRI. J. Cogn. Neurosci. 26, 606–620 (2014).

4. van Wassenhove, V., Grant, K. W. & Poeppel, D. Temporal window of integration in auditory-visual speech perception. Neuropsychologia 45, 598–607 (2007).

5. Baart, M., Lindborg, A. & Andersen, T. S. Electrophysiological evidence for differences between fusion and combination illusions in audiovisual speech perception. Eur. J. Neurosci. 46, 2578–2583 (2017).

6. Hickok, G. & Poeppel, D. The cortical organization of speech processing. Nat. Rev. Neurosci. 8, 393–402 (2007).

7. Bernstein, L. E. & Liebenthal, E. Neural pathways for visual speech perception. Front. Neurosci. 8, 386 (2014).

8. Malfait, N. et al. Different Neural Networks Are Involved in Audiovisual Speech Perception Depending on the Context. J. Cogn. Neurosci. 26, 1572–1586 (2014).

9. Hertrich, I., Dietrich, S. & Ackermann, H. Cross-modal interactions during perception of audiovisual speech and nonspeech signals: An fMRI study. J. Cogn. Neurosci. 23, 221–237 (2011).

10. Flinker, A. et al. Redefining the role of Broca’s area in speech. Proc. Natl. Acad. Sci. 112, 2871–2875 (2015).

11. Miozzo, M., Williams, A. C., McKhann, G. M. & Hamberger, M. J. Topographical gradients of semantics and phonology revealed by temporal lobe stimulation. Hum. Brain Mapp. 38, 688–703 (2017).

12. Beauchamp, M. S., Nath, A. R. & Pasalar, S. fMRI-Guided Transcranial Magnetic Stimulation Reveals That the Superior Temporal Sulcus Is a Cortical Locus of the McGurk Effect. J. Neurosci. 30, 2414–2417 (2010).

13. Szycik, G. R., Stadler, J., Tempelmann, C. & Münte, T. F. Examining the McGurk illusion using high-field 7 Tesla functional MRI. Front. Hum. Neurosci. 6, 95 (2012).

14. Beauchamp, M. S., Argall, B. D., Bodurka, J., Duyn, J. H. & Martin, A. Unraveling multisensory integration: Patchy organization within human STS multisensory cortex. Nat. Neurosci. 7, 1190–1192 (2004).

15. Nath, A. R. & Beauchamp, M. S. A neural basis for interindividual differences in the McGurk effect, a multisensory speech illusion. Neuroimage 59, 781–787 (2012).

16. Venezia, J. H., et al. Auditory, Visual and Audiovisual Speech Processing Streams in Superior Temporal Sulcus. Front. Hum. Neurosci. 11, 174 (2017).

17. Hickok, G. et al. Neural Networks Supporting Audiovisual Integration for Speech: A Large-Scale Lesion Study. Cortex 103, 360–371 (2018).

18. Bhat, J., Miller, L. M., Pitt, M. A. & Shahin, A. J. Putative mechanisms mediating tolerance for audiovisual stimulus onset asynchrony. J. Neurophysiol. 113, 1437– 1450 (2015).

19. Schwartz, J.-L. & Savariaux, C. No, There Is No 150 ms Lead of Visual Speech on Auditory Speech, but a Range of Audiovisual Asynchronies Varying from Small Audio Lead to Large Audio Lag. PLoS Comput. Biol. 10, e1003743 (2014).

20. Macaluso, E., George, N., Dolan, R., Spence, C. & Driver, J. Spatial and temporal factors during processing of audiovisual speech: A PET study. Neuroimage 21, 725– 732 (2004).

21. Olson, I. R., Gatenby, J. C. & Gore, J. C. A comparison of bound and unbound audio-visual information processing in the human cerebral cortex. *Cogn*. Brain Res. 14, 129–138 (2002).

22. Simon, D. M., Nidiffer, A. R. & Wallace, M. T. Single Trial Plasticity in Evidence Accumulation Underlies Rapid Recalibration to Asynchronous Audiovisual Speech. Sci. Rep. 8, 12499 (2018).

23. Foss-feig, J. H. et al. An extended multisensory temporal binding window in autism spectrum disorders. Exp. brain Res. 203, 381–389 (2010).

24. Coull, J. T. & Nobre, A. C. Dissociating explicit timing from temporal expectation with fMRI. Curr. Opin. Neurobiol. 18, 137–144 (2008).

25. Hironaga, N. et al. Spatiotemporal brain dynamics of auditory temporal assimilation. Sci. Rep. 7, 4–9 (2017).

26. Baumann, O. et al. Neural Correlates of Temporal Complexity and Synchrony during Audiovisual Correspondence Detection. Eneuro 5, a0294–17.2018 (2018).

27. Hagoort, P. Nodes and networks in the neural architecture for language: Broca’s region and beyond. Curr. Opin. Neurobiol. 28, 136–141 (2014).

28. Miller, L. M. & D’Esposito, M. Perceptual Fusion and Stimulus Coincidence in the Cross-Modal Integration of Speech. J. Neurosci. 25, 5884–5893 (2005).

29. Arnal, L. H., Wyart, V. & Giraud, A.-L. Transitions in neural oscillations reflect prediction errors generated in audiovisual speech. Nat. Neurosci. 14, 797–801 (2011).

30. Genovesio, A., Tsujimoto, S. & Wise, S. P. Feature- and Order-Based Timing Representations in the Frontal Cortex. Neuron 63, 254–266 (2009).

31. Charles, D. P., Gaffan, D. & Buckley, M. J. Impaired Recency Judgments and Intact Novelty Judgments after Fornix Transection in Monkeys. J. Neurosci. 24, 2037–2044 (2004).

32. Olasagasti, I., Bouton, S. & Giraud, A.-L. Prediction across sensory modalities: A neurocomputational model of the McGurk effect. Cortex 68, 61–75 (2015).

33. Alsius, A., Navarra, J., Campbell, R. & Soto-Faraco, S. Audiovisual integration of speech falters under high attention demands. Curr. Biol. 15, 839–843 (2005).

34. Colin, C., Radeau, M., Deltenre, P., Demolin, D. & Soquet, A. The role of sound intensity and stop-consonant voicing on McGurk fusions and combinations. Eur. J. Cogn. Psychol. 14, 475–491 (2002).

35. Cathiard, M.-A., Schwartz, J.-L. & Abry, C. Asking a naive question to the McGurk effect: why does audio [b] give more [d] percepts with visual [g] than with visual [d]? in AVSP 2001 International Conference on Auditory-Visual Speech Processing 138–142 (2001).

36. Soto-Faraco, S. & Alsius, A. Deconstructing the McGurk-MacDonald Illusion. J. Exp. Psychol. Hum. Percept. Perform. 35, 580–587 (2009).

37. Morís Fernández, L., Torralba, M. & Soto-Faraco, S. Theta oscillations reflect conflict processing in the perception of the McGurk illusion. Eur. J. Neurosci. 1–12 (2018). doi:10.1111/ejn.13804

38. Giordano, B. L. et al. Contributions of local speech encoding and functional connectivity to audio-visual speech perception. Elife 6, e24763 (2017).

39. Park, H., Kayser, C., Thut, G. & Gross, J. Lip movements entrain the observers’ low-frequency brain oscillations to facilitate speech intelligibility. Elife 5, e14521 (2016).

40. Kayser, S. J., Ince, R. A. A., Gross, J. & Kayser, C. Irregular speech rate dissociates auditory cortical entrainment, evoked responses, and frontal alpha. J. Neurosci. 35, 14691–14701 (2015).

41. Alm, M. & Behne, D. Audio-visual speech experience with age influences perceived audio-visual asynchrony in speech. 134, (2013).

42. van Wassenhove, V., Grant, K. W. & Poeppel, D. Visual speech speeds up the neural processing of auditory speech. Proc. Natl. Acad. Sci. U. S. A. 102, 1181–6 (2005).

43. Morís Fernández, L., Visser, M., Ventura-Campos, N., Ávila, C. & Soto-Faraco, S. Top-down attention regulates the neural expression of audiovisual integration. Neuroimage 119, 272–285 (2015).

44. Arnal, L. H., Morillon, B., Kell, C. a & Giraud, A.-L. Dual neural routing of visual facilitation in speech processing. J. Neurosci. 29, 13445–53 (2009).

45. Olasagasti, I. & Giraud, A.-L. Integrating prediction errors at two time scales permits rapid recalibration of speech sound categories. BioRxiv 1–19 (2018).

46. Auksztulewicz, R. et al. The Cumulative Effects of Predictability on Synaptic Gain in the Auditory Processing Stream. J. Neurosci. 37, 6751–6760 (2017).

47. Phillips, H. N. et al. Convergent evidence for hierarchical prediction networks from human electrocorticography and magnetoencephalography. Cortex 82, 192–205 (2016).

48. Block, R. A. & Gruber, R. P. Time perception, attention, and memory: A selective review. Acta Psychol. (Amst). 149, 129–133 (2014).

49. Kocagoncu, E., Clarke, A., Devereux, B. J. & Tyler, L. K. Decoding the Cortical Dynamics of Sound-Meaning Mapping. J. Neurosci. 37, 1312–1319 (2017).

50. Klucharev, V., Möttönen, R. & Sams, M. Electrophysiological indicators of phonetic and non-phonetic multisensory interactions during audiovisual speech perception. *Cogn*. Brain Res. 18, 65–75 (2003).

51. Adank, P., Nuttall, H., Bekkering, H. & Maegherman, G. Effects of stimulus response compatibility on covert imitation of vowels. *Attention, Perception*, Psychophys. 1–10 (2018). doi:10.3758/s13414-018-1501-3

52. Green, K. P. & Kuhl, P. K. The interaction of visual place and auditory voicing information during phonetic perception. J. Exp. Psychol. Hum. Percept. Perform. 17, 278–288 (1991).

53. Hessler, D., Jonkers, R., Stowe, L. & Bastiaanse, R. The whole is more than the sum of its parts - Audiovisual processing of phonemes investigated with ERPs. Brain Lang. 124, 213–224 (2013).

54. Keane, B. P., Rosenthal, O., Chun, N. H. & Shams, L. Audiovisual integration in high functioning adults with autism. Res. Autism Spectr. Disord. 4, 276–289 (2010).

55. Nahorna, O., Berthommier, F. & Schwartz, J.-L. Binding and unbinding the auditory and visual streams in the McGurk effect. J. Acoust. Soc. Am. 132, 1061–1077 (2012).

56. Norrix, L. W., Plante, E. & Vance, R. Auditory-visual speech integration by adults with and without language-learning disabilities. J. Commun. Disord. 39, 22–36 (2006).

57. Zhu, L. L. & Beauchamp, M. S. Mouth and Voice : A Relationship between Visual and Auditory Preference in the Human Superior Temporal Sulcus. J. Neurosci. 37, 2697– 2708 (2017).

58. Stein, B. E. & Stanford, T. R. Multisensory integration : current issues from the perspective of the single neuron. Nat. Rev. Neurosci. 9, 255–66 (2008).

59. Barraclough, N. E., Xiao, D., Baker, C. I., Oram, M. W. & Perrett, D. I. Integration of Visual and Auditory Information by Superior Temporal Sulcus Neurons Responsive to the Sight of Actions. J. Cogn. Neurosci. 17, 377–391 (2005).

60. Willems, R. M., Özyürek, A. & Hagoort, P. Differential roles for left inferior frontal and superior temporal cortex in multimodal integration of action and language. Neuroimage 47, 1992–2004 (2009).

61. Özyürek, A. Hearing and seeing meaning in speech and gesture : insights from brain and behaviour. Philos. Trans. R. Soc. B Biol. Sci. 369, 20130296 (2014).

62. Pratt, H., Bleich, N. & Mittelman, N. Spatio-temporal distribution of brain activity associated with audio-visually congruent and incongruent speech and the McGurk Effect. Brain Behav. 5, 1–25 (2015).

63. Friston, K. Does predictive coding have a future? Nat. Neurosci. 21, 1019–1021 (2018).

64. Rao, R. P. N. & Ballard, D. H. Predictive coding in the visual cortex: a functional interpretation of some extra-classical receptive-field effects. Nat. Neurosci. 2, 79–87 (1999).

65. Lüttke, C. S., Ekman, M., van Gerven, M. A. J. & de Lange, F. P. McGurk illusion recalibrates subsequent auditory perception. Sci. Rep. 6, 32891 (2016).

66. Lüttke, C. S., Pérez-Bellido, A. & de Lange, F. P. Rapid recalibration of speech perception after experiencing the McGurk illusion. R. Soc. Open Sci. 5, (2018).

67. Fontes, R. et al. Time perception mechanisms at central nervous system. Neurol. Int. 8, 14–22 (2016).

68. Ivry, R. B. & Spencer, R. M. C. The neural representation of time. Curr. Opin. Neurobiol. 14, 225–232 (2004).

69. Mangels, J. A., Ivry, R. B. & Shimizu, N. Dissociable contributions of the prefrontal and neocerebellar cortex to time perception. *Cogn*. Brain Res. 7, 15–39 (1998).

70. Rao, S. M., Mayer, A. R. & Harrington, D. L. The evolution of brain activation during temporal processing. Nat. Neurosci. 4, 317–323 (2001).

71. van Wassenhove, V. & Nagarajan, S. S. Auditory Cortical Plasticity in Learning to Discriminate Modulation Rate. J. Neurosci. 27, 2663–2672 (2007).

72. Onoe, H. et al. Cortical networks recruited for time perception: A monkey positron emission tomography (PET) study. Neuroimage 13, 37–45 (2001).

73. Di Liberto, G. M., Lalor, E. C. & Millman, R. E. Causal cortical dynamics of a predictive enhancement of speech intelligibility. Neuroimage 166, 247–258 (2018).

74. Kristensen, L. B., Engberg-Pedersen, E. & Wallentin, M. Context Predicts Word Order Processing in Broca’s Region. J. Cogn. Neurosci. 26, 2762–2777 (2016).

75. Matchin, W., Hammerly, C. & Lau, E. The role of the IFG and pSTS in syntactic prediction: Evidence from a parametric study of hierarchical structure in fMRI. Cortex 88, 106–123 (2017).

76. Grabot, L., Kösem, A., Azizi, L. & Van Wassenhove, V. Prestimulus Alpha Oscillations and the Temporal Sequencing of Audio-visual Events. J. Cogn. Neurosci. 29, 168– 169 (2017).

77. Morís Fernández, L., Macaluso, E. & Soto-Faraco, S. Audiovisual integration as conflict resolution: The conflict of the McGurk illusion. Hum. Brain Mapp. 38, 5691–5705 (2017).

78. Shahin, A. J., Backer, K. C., Rosenblum, L. D. & Kerlin, J. R. Neural mechanisms underlying cross-modal phonetic encoding. J. Neurosci. 1566–17 (2017). doi:10.1523/JNEUROSCI.1566-17.2017

79. Tadel, F., Baillet, S., Mosher, J. C., Pantazis, D. & Leahy, R. M. Brainstorm: A user-friendly application for MEG/EEG analysis. Comput. Intell. Neurosci. 879716 (2011). doi:10.1155/2011/879716

80. Cointepas, Y., Geffroy, D., Souedet, N. & Denghien, I. The BrainVISA project: a shared software development infrastructure for biomedical imaging research. in Proceedings 16th HBM (2010).

81. Hämäläinen, M., Hari, R., Ilmoniemi, R. J., Knuutila, J. & Lounasmaa, O. V. Magnetoencephalography theory, instrumentation, and applications to noninvasive studies of the working human brain. Rev. Mod. Phys. 65, 413–497 (1993).

82. Meyers, E. M. The Neural Decoding Toolbox. Front. Neuroinform. 7, 8 (2013).

83. Isik, L., Meyers, E. M., Leibo, J. Z. & Poggio, T. The dynamics of invariant object recognition in the human visual system. J. Neurophysiol. 111, 91–102 (2014).

84. Boran, E. et al. Persistent hippocampal neural firing and hippocampal-cortical coupling predict verbal working memory load. Sci. Adv. 5, eaav3687 (2019).

